# Low-Cost Touchscreen Driven Programmable Dual Syringe Pump for Life Science Applications

**DOI:** 10.1101/288290

**Authors:** Valentina E. Garcia, Jamin Liu, Joseph L. DeRisi

## Abstract

Syringe pumps are powerful tools able to automate routine laboratory practices that otherwise consume large amounts of manual labor time. Commercially available syringe pumps are expensive, difficult to customize, and often preset for a narrow range of operations. Here, we show how to build a programmable dual syringe pump (PDSP) that overcomes these limitations. The PDSP is driven by a Raspberry Pi paired with a stepper motor controller to allow maximal customization via Python scripting. The entire setup can be controlled by a touchscreen for use without a keyboard or mouse. Furthermore, the PDSP is structured around 3D printed parts, enabling users to change any component for their specific application. We demonstrate one application of the PDSP by using it to generate whole cell lysates using a cell homogenizer in an automated fashion.

**Specifications table:** 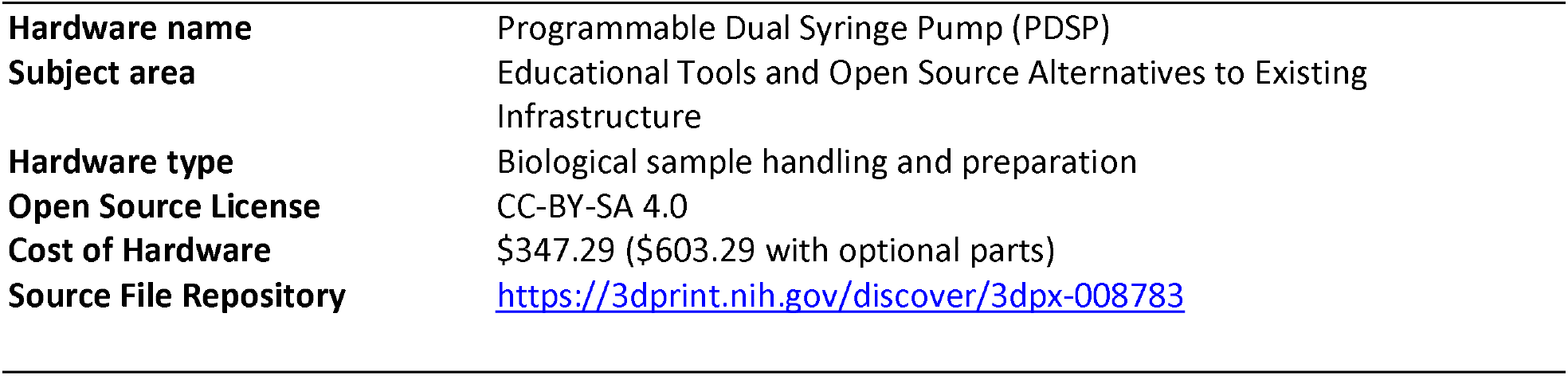

## 1. Hardware in context

Syringe pumps have a wide variety of uses across fields from engineering to biology. Their primary purpose is to continuously dispense precise volumes over a set amount of time. They save time by running unsupervised and provide more consistency than human hands. A dual syringe pump allows for two syringes to have coordinated actions, broadening the potential applications. In particular, dual syringe pumps have the power to automate a number of routine and repetitive protocols in the life sciences.

Our lab initially conceived of a dual syringe pump that could be used to make *Plasmodium falciparum* whole cell extracts used for *in vitro* translation. Previously, we generated lysates by passing purified infected red blood cells through an Isobiotech cell homogenizer using a ball bearing with 4um clearance. Frozen pellets of purified parasites were thawed, loaded into one 3mL syringe, and passed through the homogenizer into a second 3mL syringe. The lysate was then passed back into the first syringe. This back and forth cycle was repeated up to 20 times per parasite pellet^1^. This process takes between 20-30 mins per parasite pellet and the resistance in the device makes it physically taxing on the wrists and thumbs. In the lab, lysate generation was often a rate-limiting step when making *in vitro* translation extracts due to the manual and tedious nature of the process. It also entailed unacceptable amounts of user-dependent variation between lysate preparations. In the worst cases, syringes would break or the plunger would deform as pressure was applied unevenly. A customizable PDSP addresses all of these issues not only for our uses, but also for other applications of the Isobiotech Cell homogenizer such as *C. elegans* lysis^2^ and mammalian cell culture homogenization^3^.

We reasoned that this process could be easily replaced with a programmable dual syringe pump. However, commercially available dual syringe pumps have a number of limitations, including cost and flexibility. As of this writing, commercially available dual syringe pumps cost upwards of $1500^4^ and coordinated programmable motion tends to be limited. There are currently no solutions that feature a customizable graphical user interface (GUI) touchscreen, which greatly simplify use. Additionally, the physical configuration of the commercial products makes alternative mounting options difficult. To address these issues we designed and built a PDSP that can be further customized for any use.

## 2. Hardware description

Our custom PDSP is constructed on an extruded aluminum frame that can be mounted horizontally or vertically. For our specific application, we chose a vertical mount to allow the cell homogenizer to be immersed in ice (Fig. 1A/1B). The PDSP utilizes two NEMA-14 stepper motors (StepperOnline) that create precise volume changes even at high-torque. The motors are driven by a Pi-Plate MOTORplate controller (Pi-Plates, Inc.) mated to a Raspberry Pi (v3 Model B). Integrated limit switches provide for simple and accurate “homing” procedures. To make the PDSP easy to use for routine laboratory stand-alone use, the PDSP is operated via an attached touchscreen (Landzo), without a keyboard or mouse.

**Figure 1:**
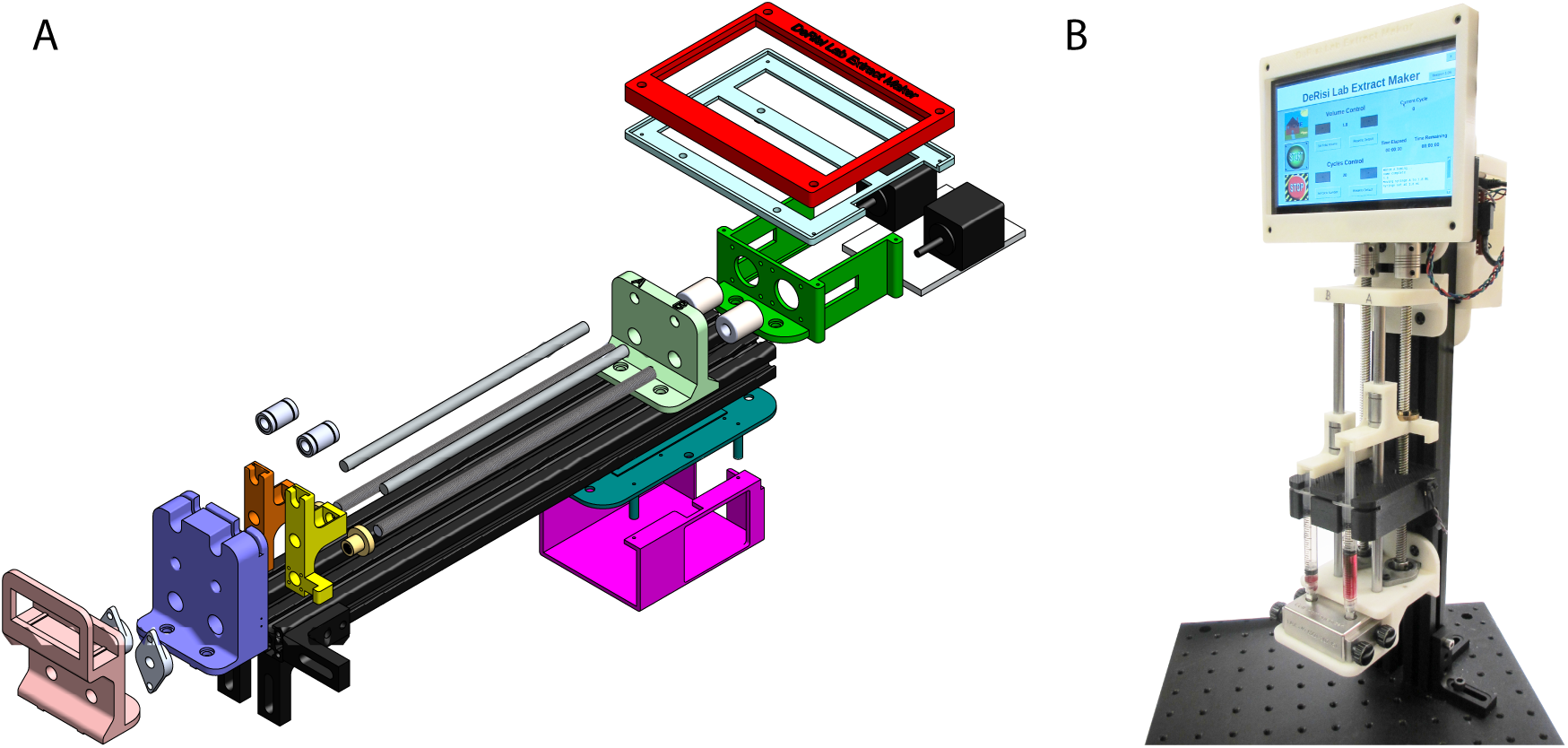
A) A 3D rendering of the PDSP with all components visible. B) A photograph of the completed PDSP standing vertically.

While this PDSP is specialized to lysate generation, alternative applications may have different requirements. We designed this device so that it could be adapted to many different environments, such as a biohazard hood, and many different tasks, such as microfluidic experiments. To allow for maximum customization, we made the PDSP modular with 3D printed parts that can be interchanged to accommodate different volume syringes. By simply changing the dimensions of the printed parts the syringes can be set to any distance apart. The cell homogenizer holder can be interchanged with any other user-designed holder. Our custom Python/Tkinter interface can be easily configured to drive the stepper motors at different speeds allowing for a range of flow rates or even gradients of flow rates. The Pi-Plate MOTORplates can be stacked, allowing a single Pi and interface to simultaneously control up to 8 PDSPs (16 syringes) for high volume production environments.

## 3. Design files

Table 1 contains links to the STL design files for all 3D printed parts described here:

**Table 1.**
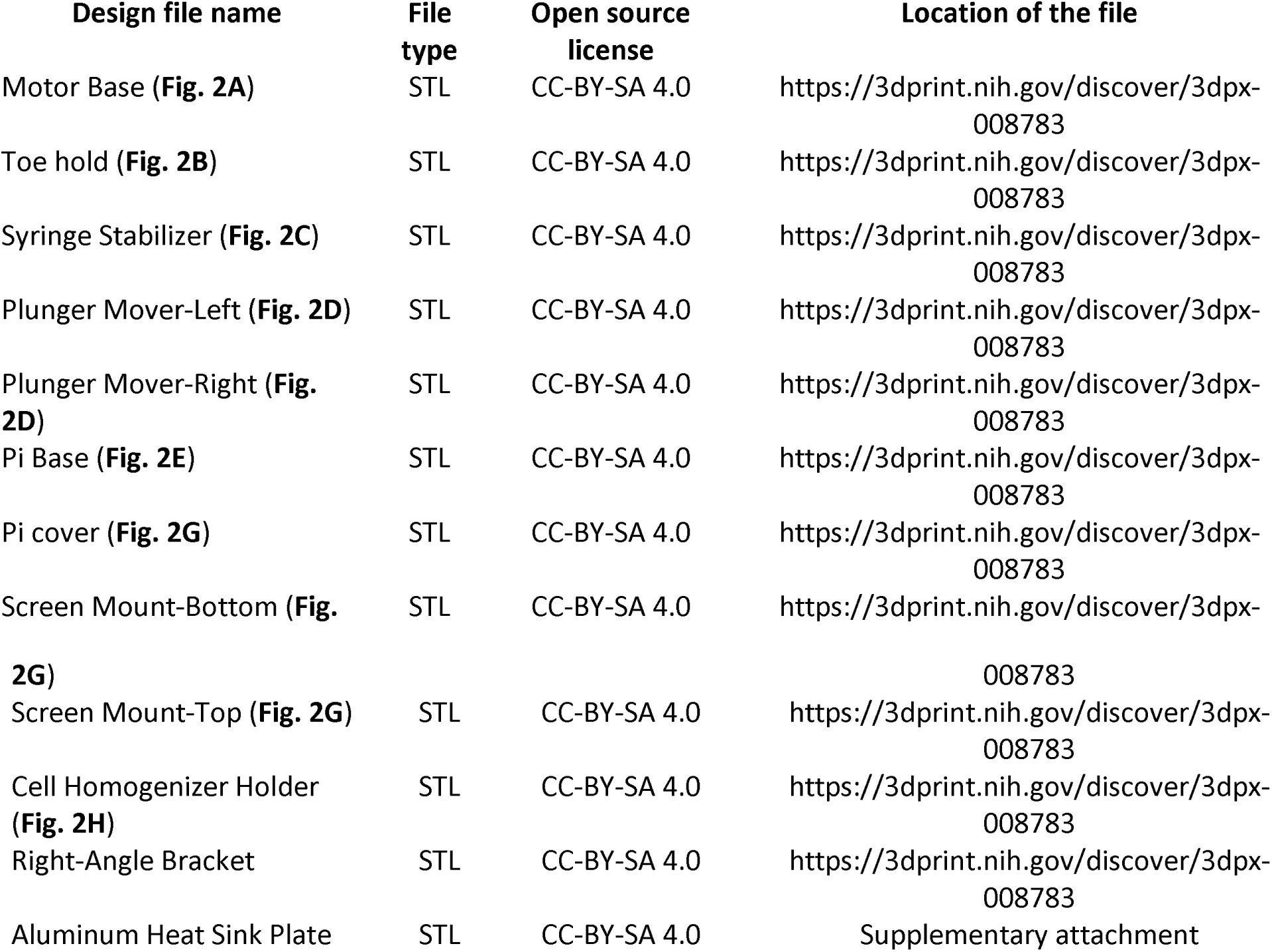
Design files for all 3D printed parts

- The **Motor Base** (Fig. 2A) secures the two stepper motors and holds the Aluminum Heat Sink Plate against the T-profile rails.

**Figure 2:**
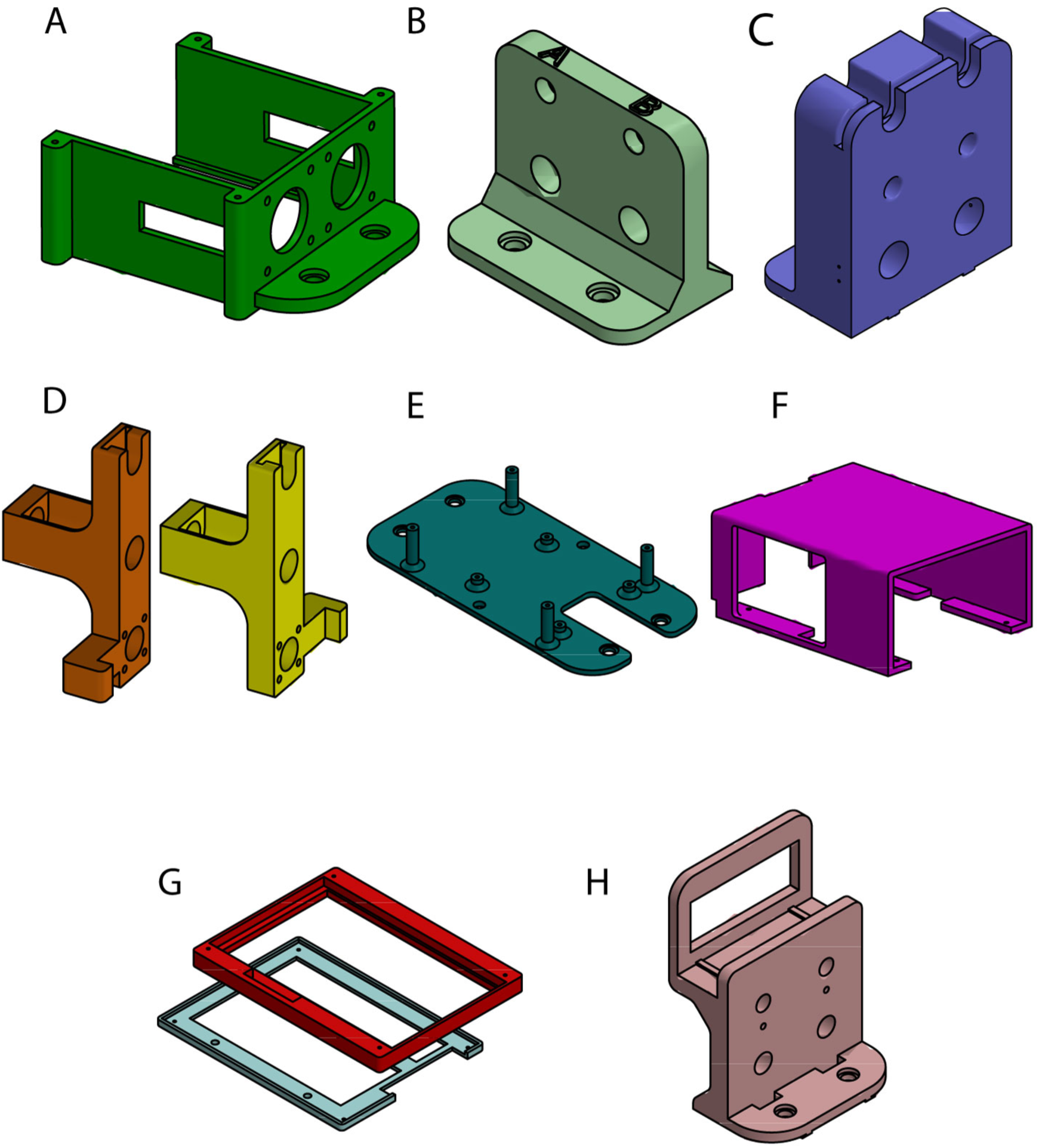
Models of all 3D printed PDSP components: A) Motor Base, B) Toe Hold, C) Syringe Stabilizer, D) Left and right Plunger Movers, E) Pi Base, F) Pi Cover, G) Top and bottom Screen Mounts, and H) Cell Homogenizer Holder
- The **Toe Hold** (Fig. 2B) positions the linear motion shafts against the T-profile rails.
- The **Syringe Stabilizer** (Fig. 2C) holds the syringe body in place by securing both the barrel and the barrel flange parallel to the movement direction of the plunger.
- The left and right **Plunger Movers** (Fig. 2D) hold the plunger flange of the syringes and move them along the linear motion shaft as the T8 threaded rods turn according to the stepper motors to change the volume.
- The **Pi Base** (Fig. 2E) and **Cover** (Fig. 2F) secure the Raspberry Pi along the T-profile rails and insulates it from short-circuiting against nearby conductive material. It also protects the Raspberry Pi from dust and accidental splashes from the ice bucket.
- The **Screen Mount Top** and **Bottom** (Fig. 2G) hold the touch screen at eye-level when the PDSP is constructed vertically. Situated in front of the motors, it keeps the touchscreen a safe distance from the generated heat.
- The **Cell Homogenizer Holder** (Fig. 2H) positions the cell homogenizer in alignment with luer lock syringes such that no additional tubing is necessary. It also allows the cell homogenizer to be in contact with ice at all times.
- The **Right-Angle Brackets** are used to position the syringe pump vertically on a breadboard. If desired, they can be 3D printed rather than purchased.
- The **Aluminum Heat Sink Plate** is placed between the stepper motors on the Motor Base and the T-profile rails to rapidly disperse heat generated from the stepper.

## 4. Bill of Materials

**Table 2.**
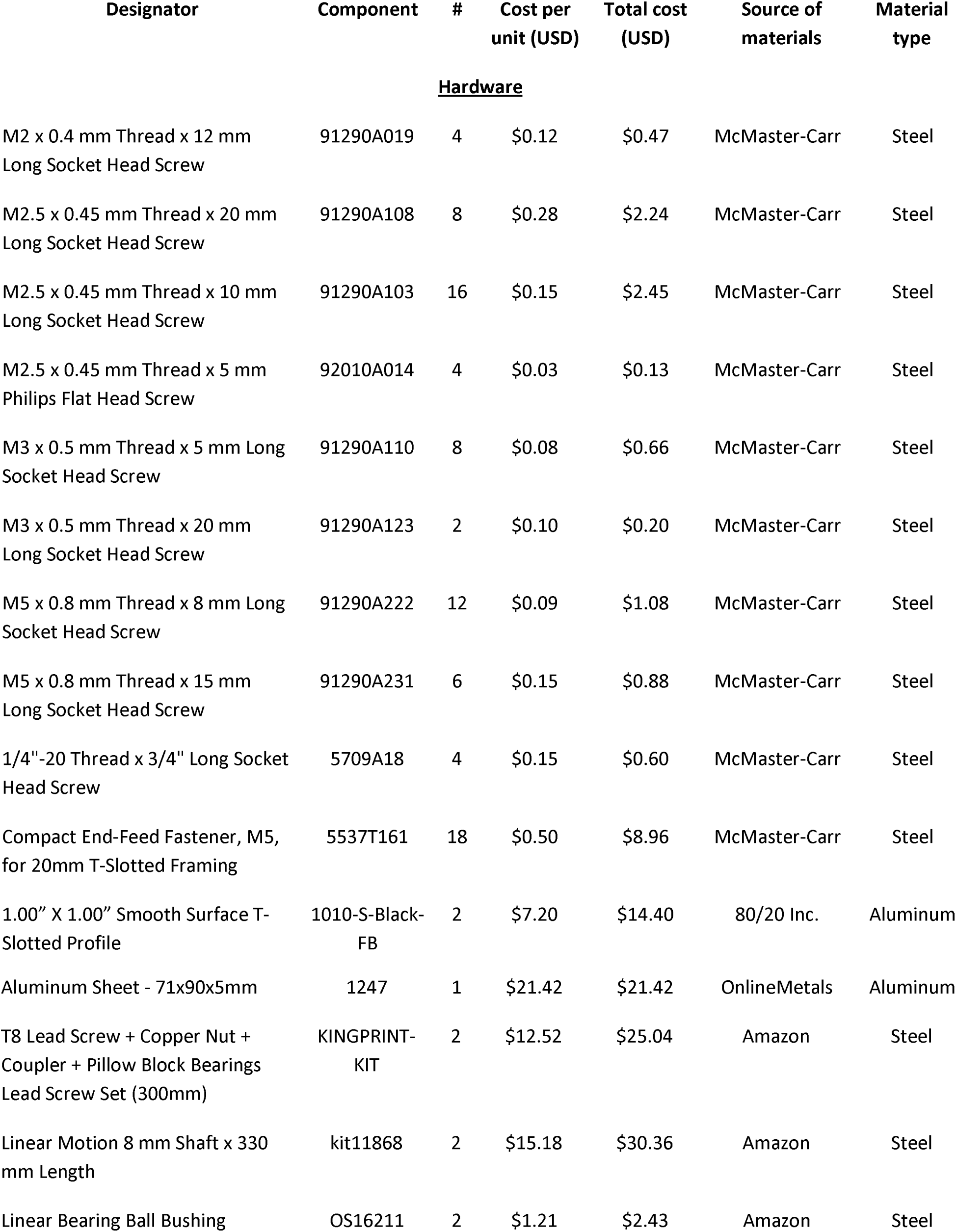
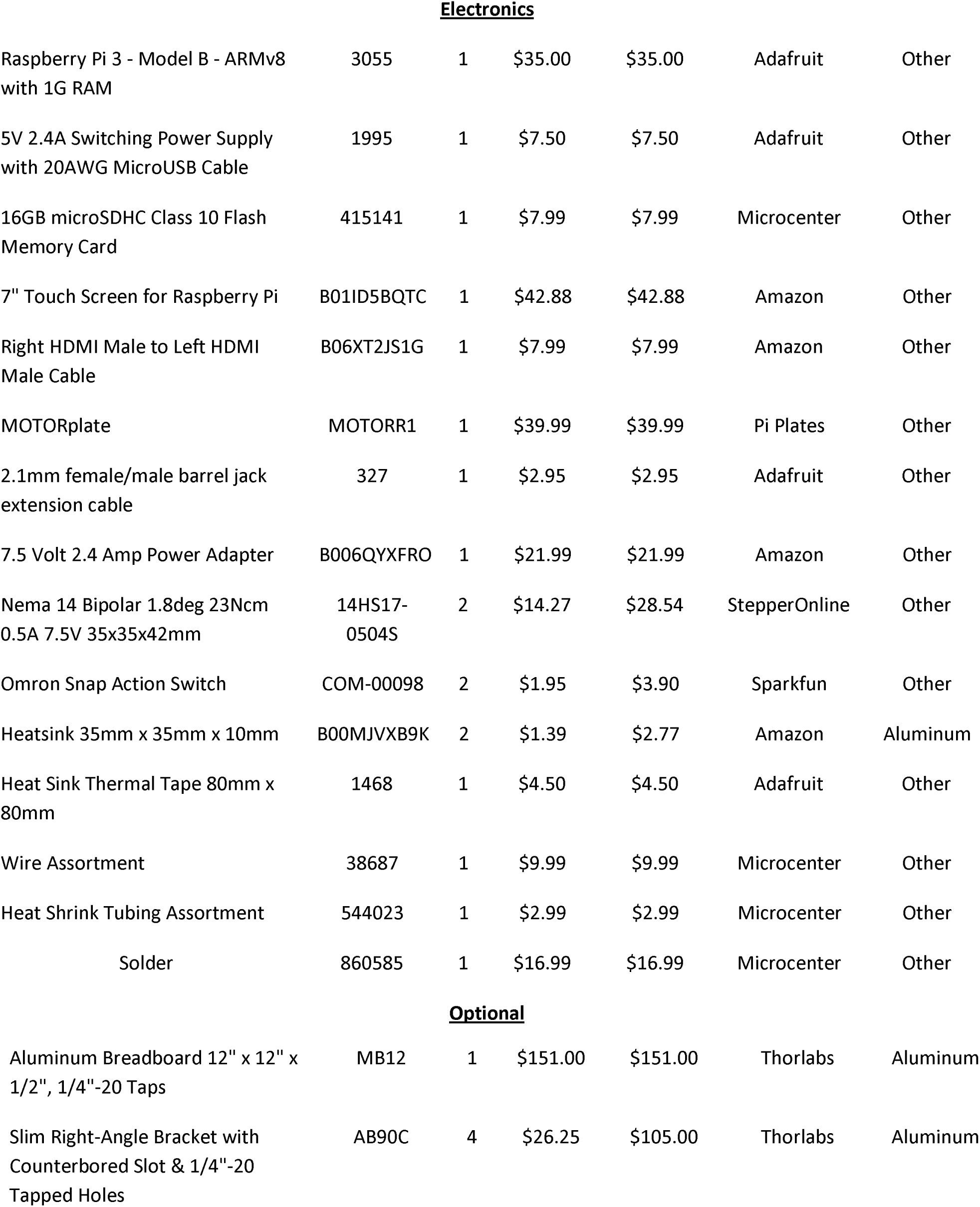
List of all purchased items with number required and cost

| Designator | Component | # | Cost per unit (USD) | Total cost (USD) | Source of materials | Material type |
| --- | --- | --- | --- | --- | --- | --- |
| **<u>Hardware</u>** |  |  |  |  |  |  |
| M2 x 0.4 mm Thread x 12 mm Long Socket Head Screw | 91290A019 | 4 | \$0.12 | \$0.47 | McMaster-Carr | Steel |
| M2.5 x 0.45 mm Thread x 20 mm Long Socket Head Screw | 91290A108 | 8 | \$0.28 | \$2.24 | McMaster-Carr | Steel |
| M2.5 x 0.45 mm Thread x 10 mm Long Socket Head Screw | 91290A103 | 16 | \$0.15 | \$2.45 | McMaster-Carr | Steel |
| M2.5 x 0.45 mm Thread x 5 mm Philips Flat Head Screw | 92010A014 | 4 | \$0.03 | \$0.13 | McMaster-Carr | Steel |
| M3 x 0.5 mm Thread x 5 mm Long Socket Head Screw | 91290A110 | 8 | \$0.08 | \$0.66 | McMaster-Carr | Steel |
| M3 x 0.5 mm Thread x 20 mm Long Socket Head Screw | 91290A123 | 2 | \$0.10 | \$0.20 | McMaster-Carr | Steel |
| M5 x 0.8 mm Thread x 8 mm Long Socket Head Screw | 91290A222 | 12 | \$0.09 | \$1.08 | McMaster-Carr | Steel |
| M5 x 0.8 mm Thread x 15 mm Long Socket Head Screw | 91290A231 | 6 | \$0.15 | \$0.88 | McMaster-Carr | Steel |
| 1/4"-20 Thread x 3/4" Long Socket Head Screw | 5709A18 | 4 | \$0.15 | \$0.60 | McMaster-Carr | Steel |
| Compact End-Feed Fastener, M5, for 20mm T-Slotted Framing | 5537T161 | 18 | \$0.50 | \$8.96 | McMaster-Carr | Steel |
| 1.00" X 1.00" Smooth Surface T-Slotted Profile | 1010-S-Black-FB | 2 | \$7.20 | \$14.40 | 80/20 Inc. | Aluminum |
| Aluminum Sheet - 71x90x5mm | 1247 | 1 | \$21.42 | \$21.42 | OnlineMetals | Aluminum |
| T8 Lead Screw + Copper Nut + Coupler + Pillow Block Bearings Lead Screw Set (300mm) | KINGPRINT-KIT | 2 | \$12.52 | \$25.04 | Amazon | Steel |
| Linear Motion 8 mm Shaft x 330 mm Length | kit11868 | 2 | \$15.18 | \$30.36 | Amazon | Steel |
| Linear Bearing Ball Bushing | OS16211 | 2 | \$1.21 | \$2.43 | Amazon | Steel |
| **Electronics** |  |  |  |  |  |  |
| Raspberry Pi 3 - Model B - ARMv8 with 1G RAM | 3055 | 1 | \$35.00 | \$35.00 | Adafruit | Other |
| 5V 2.4A Switching Power Supply with 20AWG MicroUSB Cable | 1995 | 1 | \$7.50 | \$7.50 | Adafruit | Other |
| 16GB microSDHC Class 10 Flash Memory Card | 415141 | 1 | \$7.99 | \$7.99 | Microcenter | Other |
| 7" Touch Screen for Raspberry Pi | B01ID5BQTC | 1 | \$42.88 | \$42.88 | Amazon | Other |
| Right HDMI Male to Left HDMI Male Cable | B06XT2JS1G | 1 | \$7.99 | \$7.99 | Amazon | Other |
| MOTORplate | MOTORR1 | 1 | \$39.99 | \$39.99 | Pi Plates | Other |
| 2.1mm female/male barrel jack extension cable | 327 | 1 | \$2.95 | \$2.95 | Adafruit | Other |
| 7.5 Volt 2.4 Amp Power Adapter | B006QYXFRO | 1 | \$21.99 | \$21.99 | Amazon | Other |
| Nema 14 Bipolar 1.8deg 23Ncm 0.5A 7.5V 35x35x42mm | 14HS17-0504S | 2 | \$14.27 | \$28.54 | StepperOnline | Other |
| Omron Snap Action Switch | COM-00098 | 2 | \$1.95 | \$3.90 | Sparkfun | Other |
| Heatsink 35mm x 35mm x 10mm | B00MJVXB9K | 2 | \$1.39 | \$2.77 | Amazon | Aluminum |
| Heat Sink Thermal Tape 80mm x 80mm | 1468 | 1 | \$4.50 | \$4.50 | Adafruit | Other |
| Wire Assortment | 38687 | 1 | \$9.99 | \$9.99 | Microcenter | Other |
| Heat Shrink Tubing Assortment | 544023 | 1 | \$2.99 | \$2.99 | Microcenter | Other |
| Solder | 860585 | 1 | \$16.99 | \$16.99 | Microcenter | Other |
| **Optional** |  |  |  |  |  |  |
| Aluminum Breadboard 12" x 12" x 1/2", 1/4"-20 Taps | MB12 | 1 | \$151.00 | \$151.00 | Thorlabs | Aluminum |
| Slim Right-Angle Bracket with Counterbored Slot & 1/4"-20 Tapped Holes | AB90C | 4 | \$26.25 | \$105.00 | Thorlabs | Aluminum |

## 5. Build Instructions

The essential PDSP components are 3D printed. The remaining hardware can be acquired online from common hardware suppliers, such as McMaster-Carr. All electronic parts can be acquired from common electronics suppliers, such as Adafruit or Microcenter (see **Bill of Materials**).

The additional tools required for assembly include:

- M2.5 Tap
- M3 Tap
- Metric Hex Key Set
- Imperial Hex Key Set
- Philips Screwdriver
- Wire Stripper
- Soldering iron
- Hack-saw or chop-saw
- Drill Press

### 5.1 Preparation of electronic components

It is useful to prepare the electronic components prior to assembly. First, solder wires to the GND pin and the N pin on the leaf switches as outlined in Fig. 3. Wrap the exposed solder and pin with heat shrink to protect the connection. To keep the electronic wiring neat, braid the four wires of each stepper motor and twist together the two wires of each leaf switches. Next, attach one heat sink onto the rear of each stepper motor by applying an appropriately sized piece of thermal tape to the rear end-cap and firmly pressing the heat sink to the tape. To prepare the motor power supply, cut and strip the wires of the 2.1mm female/male barrel jack extension cable to separate the power and ground wires.

**Figure 3:**
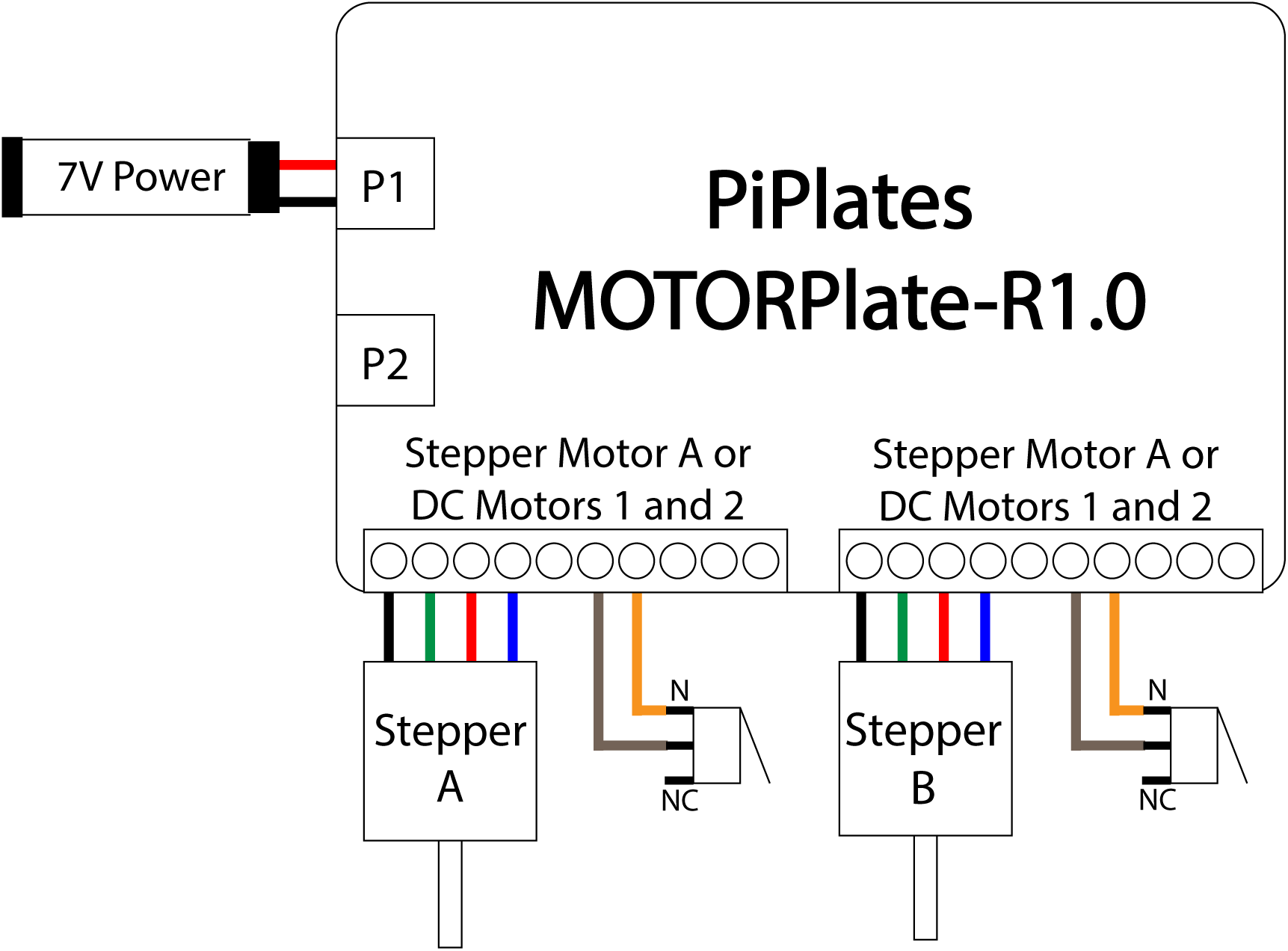
Electrical wiring schematic of the PDSP

Finally, attach the Raspberry Pi onto the Pi base (Fig. 2E) using four M2.5×5mm screws. Next, attach the PiPlates Motor Plate to the Raspberry Pi header pins by carefully applying even pressure on the plate while pushing the two components together to prevent bending any pins. Once attached, secure the PiPlates Motor Plate onto the Pi base using another four M2.5 × 5mm screws.

### 5.2 Hardware Preparation

3D print each of the necessary parts from the provided design files, including four right-angle brackets if not using ThorLab’s precision cut components. We recommend printing with ABS material with a low-density fill. Cut the two T8 lead screws to 25cm each and the two linear motion shafts to 21.5cm each. Saw the aluminum plate to the appropriate dimensions according the Aluminum Heat Sink Plate STL file (**Supplementary File 1**). Using a drill press, drill two through holes as specified into the aluminum plate for later assembly onto the T-profile rails. Tap M2.5 and M3 holes into the 3D printed parts as indicated in their respective STL files.

### 5.3 Hardware Assembly

The complete assembly of the PDSP as described below can be seen as a time-lapse in **Supplementary file 2**.

1. Attach the compact end-feed fastener with M5 x 5mm screws to each through hole of the Motor Base (Fig. 2A), Toe Hold (Fig. 2B), Syringe Stabilizer (Fig. 2C), Pi Base (Fig. 2E), and Cell Homogenizer Holder (Fig. 2H). Attach the compact end-fasteners of two M5 × 8mm through the previously drilled holes aluminum Heat Sink Plate (Fig. 4A).

**Figure 4:**
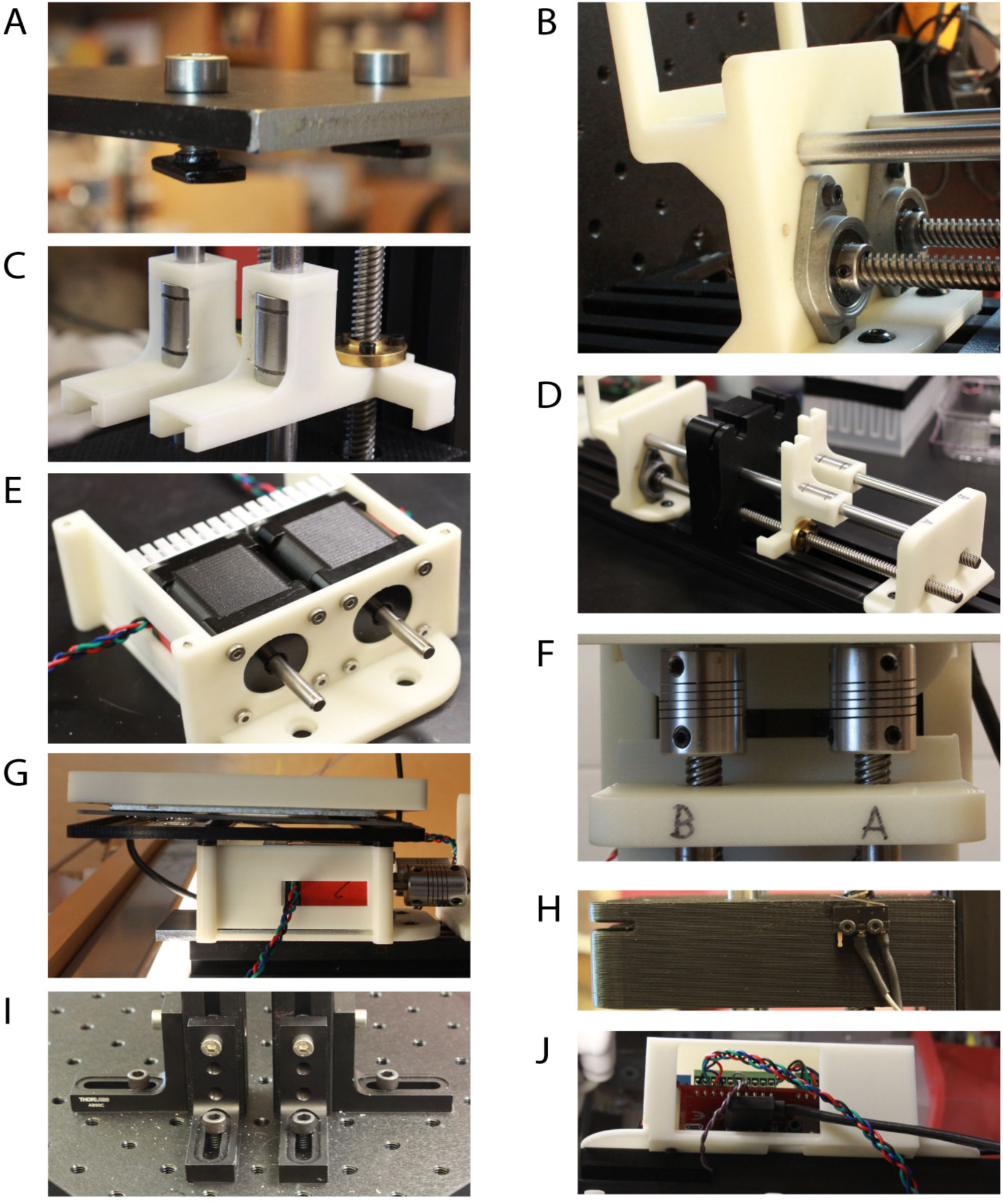
Images of key PDSP assembly steps as described in 5.4 Hardware Assembly. A) Example of the compact end-feed fastener assembly. B) The pillow block bearings, the threaded rods, and the smooth rods attached to the Extract Maker Holder. C) the copper nut and linear motion shaft assembled with the Syringe Movers on the smooth and threaded rods. D) The Toe Hold, the Syringe Holder, and the Cell Homogenizer Holder assembled on the extruded rails. E) The two stepper motors assembled with the Motor Base. F) The couplers attached to each motor and the threaded rods. G) The Screen Mount Top and Bottom sandwiching the touchscreen and assembled on the Motor Base. H) The limit switches attached to the Syringe Stabilizer component. I) The Right-Angle Brackets positioned on the aluminum breadboard to stand the PDSP vertically. J) The Raspberry Pi and PiPlates Motor Plate attached to the Pi base.
2. Attach each of the two pillow block bearings to the Extract Maker Holder (Fig. 4B) using M3 × 20mm socket head screws.
3. Insert one copper nut and one linear ball bearing into each of the Plunger Movers (Fig. 2D). Attach the copper nut using four M2.5 × 5mm socket head screws per nut and secure the linear ball bearing into its designated slot with silicone if necessary (Fig. 4C).
4. Thread the one T8 lead screws into the copper nut and slide one linear motion shaft through the linear ball bearing on each of the two Plunger Movers (Fig. 4C).
5. Making sure each part is oriented correctly, slide the four metal bars through the Syringe Stabilizer, and secure into the Cell Homogenizer Holder. The arms on each of the plunger mover should be on the outside of the linear structure and each pointed towards the Syringe Stabilizer (Fig. 4B and 4D).
6. On the other end, slide the four metal bars towards the Toe Hold making sure the linear motion shaft is secured by the Cell Homogenizer Holder on one end and by the Toe Hold on the other end (Fig. 4D).
7. Slide the Toe Hold, the Syringe Holder, and the Cell Homogenizer Holder together onto the T-profile rails by correctly slotting the compact end-feeder into the rails. Do not tighten the M5 screws yet (Fig. 4D).
8. Attach and secure the stepper motors onto the motor base using four M3 x 5mm per motor (Fig. 4E).
9. Slide the motor base with attached motors onto the T-profile rails on the side closest to the Toe Hold. Do not yet tighten the M5 screws.
10. Attach the couplers between the T8 threaded rods and the stepper motors by carefully sliding all the moving parts along the T-profile rails towards each other. After making sure that the T8 threaded rods and the motors are completely aligned, secure the couplers by tightening the attached screws (Fig. 4F).
11. Slide the Aluminum Heat Sink Plate onto the T-profile rails beneath the stepper motors on the Motor Base.
12. If desired, ensure there is sufficient space between the Cell Homogenizer Holder and the end of the T-profile rails to place an ice bucket. Finally, secure all the parts to the T-profile rails by tightening all ten of the M5 screws.
13. Attach the Screen Mount-Bottom (Fig. 2I) onto the Motor Base using four M2.5 x 20mm screws. The correct orientation should allow the touchscreen to rest on the base without bending the ribbon cable. Then, sandwich the touchscreen between the Screen Mounts using four more M2.5 x 20mm screws (Fig. 4G).
14. Attach the leaf switches associated with each stepper motor to the Syringe Stabilizer on their respective sides using M2 x 12mm screws (Fig. 4H).
15. To stand the PDSP vertically, insert one M5 x 8mm screw into the highest through hole on each of the four Right-Angle Brackets. Attach the compact end-feeder into each of these screws and slide them into the bottom of the T-profile rails, closest to the cell homogenizer holder. Do this for the four Right-Angle Brackets for a total of one each on the left and right, and two in the rear (Fig. 3I).
16. Stand the PDSP up over the breadboard and secure the Right-Angle Brackets onto the base of choice using the ¼” × ¾” long socket screws.
17. Attach the Pi Base on the rear of the PSDP such that it is flush against the top of the T-profile rails (Fig. 4I)
18. Thread the wires neatly using the through holes on the Pi Base before connecting each to the Raspberry Pi according to the Electronic Wiring Schematics (Fig. 3).
19. Slide on and secure the Pi Cover (Fig. 2H) to the rear of the Pi Base using M2.5 x 10mm screws (Fig. 4I).
20. Check that all the screws on the PDSP are appropriately tightened prior to use.

### 5.4 Software setup

To install our software and GUI, we have provided the Raspberry Pi image on our lab website (http://derisilab.ucsf.edu/). Download the image onto a MicroSD card. We suggest using the free software Etcher to create bootable SD cards^5^. Once installed, unmount the MicroSD and insert it into the Raspberry Pi. Plug in the Raspberry Pi power cord to a conventional outlet to turn it on. To power the stepper motors, connect a conventional 7.5V power supply to the barrel jack extension cable attached to the PiPlate Motor Plate.

Alternatively, install the latest release of Raspbian for the Raspberry Pi, the MOTORplate drivers (via the PIP Python repository: sudo pip install pi-plates), and the source code for the PDSP (**Supplementary File 3**) and interface manually. The resolution of the Pi can be set to fit any screen by altering the /boot/config.txt file. For the touchscreen used here, add the following lines to the end of the config.txt file:

~~~
max_usb_current=1
hdmi_group=2
hdmi_mode=1
hdmi_mode=87
hdmi_cvt 800 480 60 6 0 0 0
~~~

## 6. Operation Instructions

We have designed a GUI optimized for safe and streamlined operation of the PDSP as an automated cell homogenizer (Fig. 5). The software was written with internal limitations to protect both the sample being lysed and the PDSP components. For the push-pull pumping motion, we strongly encourage users to consider employing our intuitive GUI to operate the PDSP. The complete operation instructions of the PDSP as an automated cell homogenizer is described below and can be seen as a video clip in **Supplementary File 2**.

**Figure 5:**
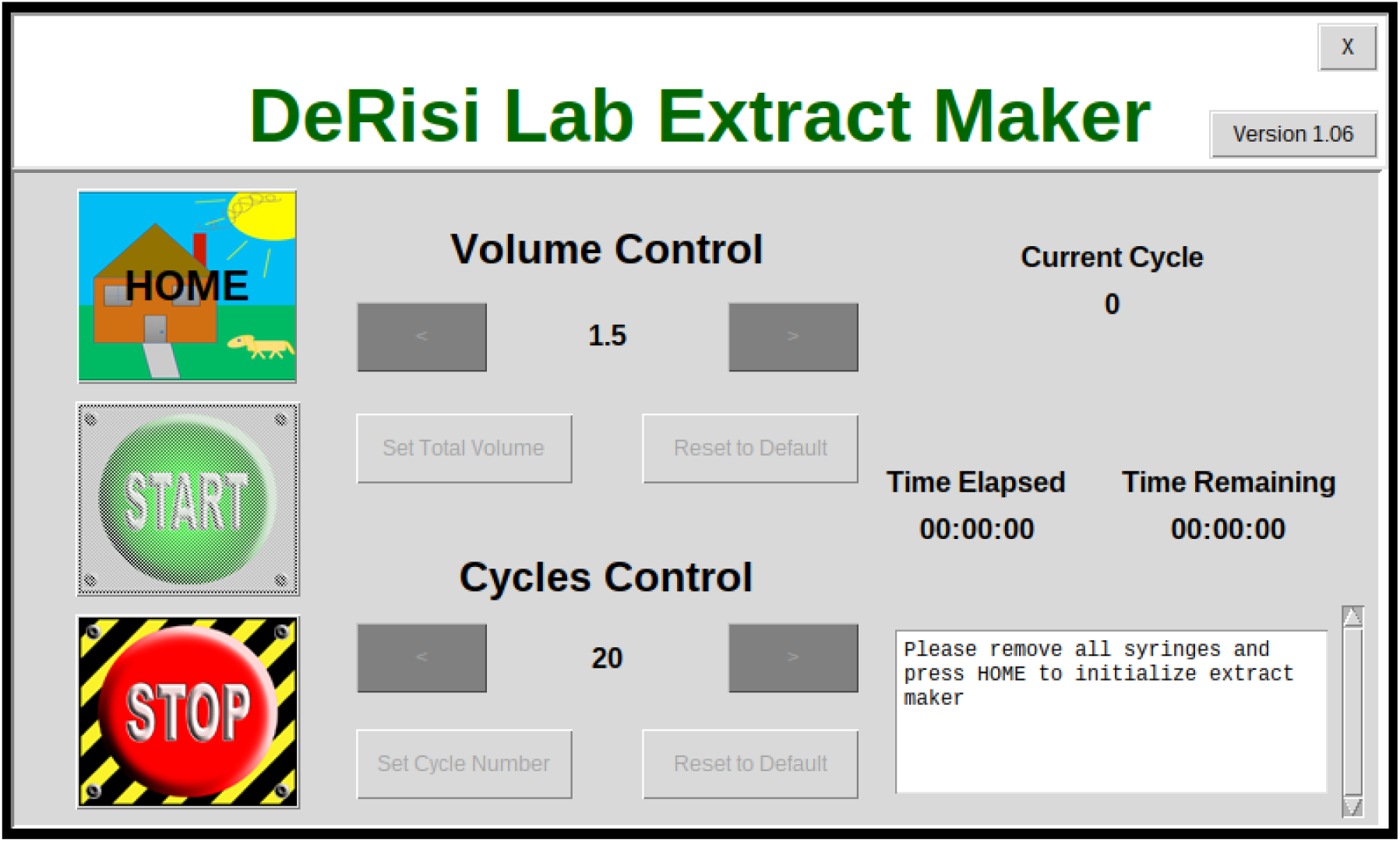
GUI interface designed in Python using Tkinter for use of the PDSP with the cell homogenizer

### 6.1 Operation Instructions for the provided GUI

1. Turn on the Raspberry Pi by plugging it in. Once the Desktop is loaded, the terminal application will launch and open the provided GUI in full-screen automatically (Fig. 5).
2. Once the GUI has launched, press the “*HOME*” button to bring both Plunger Movers to their starting position against the Syringe Stabilizer. Pay attention to the popup alerts and ensure that the PDSP is empty before homing.
3. Load a minimum of 1mL of sample into the cell homogenizer as usual. Make sure the sample is in only one syringe before loading it onto the PDSP.
4. Using the left and right arrow keys under “Volume Control” set the volume of sample in the loaded syringe. Once again, follow the popup alert instructions and ensure the cell homogenizer and syringes have not yet be placed onto the PDSP.
5. Place a filled ice bucket underneath the Cell Homogenizer Holder and insert the cell homogenizer onto the device. Ensure that each syringe is held appropriately by both the Syringe Stabilizer and the left or right Plunger Movers.
6. Using the left and right arrow keys under “Cycle Control,” set the desired number of push-pull cycles. If the cycle number is not explicitly set, the software will default to 20 cycles.
7. Press “*START*.” The number of elapsed cycles along with two timers, a countdown timer based on the estimated duration and a time elapsed counter, should appear.
8. When in doubt, there is an emergency “*STOP*” button that will immediately stop and reset all motors. To reinitialize, remove the cell homogenizer and syringes from the PDSP. Then, return to step 2 outlined here.

### 6.2 Customization Suggestions for Operation

While we use the PDSP to automate *P. falciparum* lysis using an Isobiotech Cell Homogenizer, the PDSP is adaptable to other tasks. As previously described, the PDSP is programmed on and executed from a Raspberry Pi running the Raspbian operating system. Our software to control the PDSP is written in Python using commands from the PiPlates MOTORPlate Users Guide documentation^6^. Our object oriented GUI is also designed in Python using Tkinter.

To customize the PDSP for a variety of other applications, users can reference both the script that we have written as well the extensive documentation provided by PiPlate. The PiPlate MOTORplate is highly customizable, offering a wide variety of options for stepper movement in terms of stepper size, speed, and acceleration or deceleration. Furthermore, the PiPlates MOTOR library is highly compatible with the GPIO control library, allowing flexibility in the control of the stepper motors. While python scripts can be executed from the Raspberry Pi command line using a keyboard, users can also create a specialized GUI for their own purposes.

The PDSP presented here is already compatible with a number of different pumping systems. These include, but are not limited to, continuous infusion systems, dual injection systems, and inverse linear constant flow systems. While this PDSP is relatively simple, we can add complexity by attaching up to eight PiPlate MOTORplates powered by one Raspberry Pi, allowing for control of up to sixteen stepper motors simultaneously.

## 7. Validation and Characterization

### 7.1. Characterization of the PDSP as a syringe pump

To characterize the PDSP, we tested the minimum and maximum flow rates provided by the stepper motors on a 3 mL syringe. By timing the flow of liquid, we were able to measure the consistency and the dynamic range of flow rates for the PDSP (Fig. 6A). We compared these values with the theoretical minimum and maximum flow rates that were calculated as described below:

Let the syringe cross-sectional area have units of mm^2^ and be defined as:

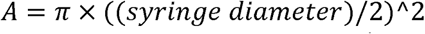

Let the linear distance conversion factor (C) have units of mm/degree and be defined as:

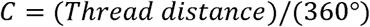

**Figure 6.**
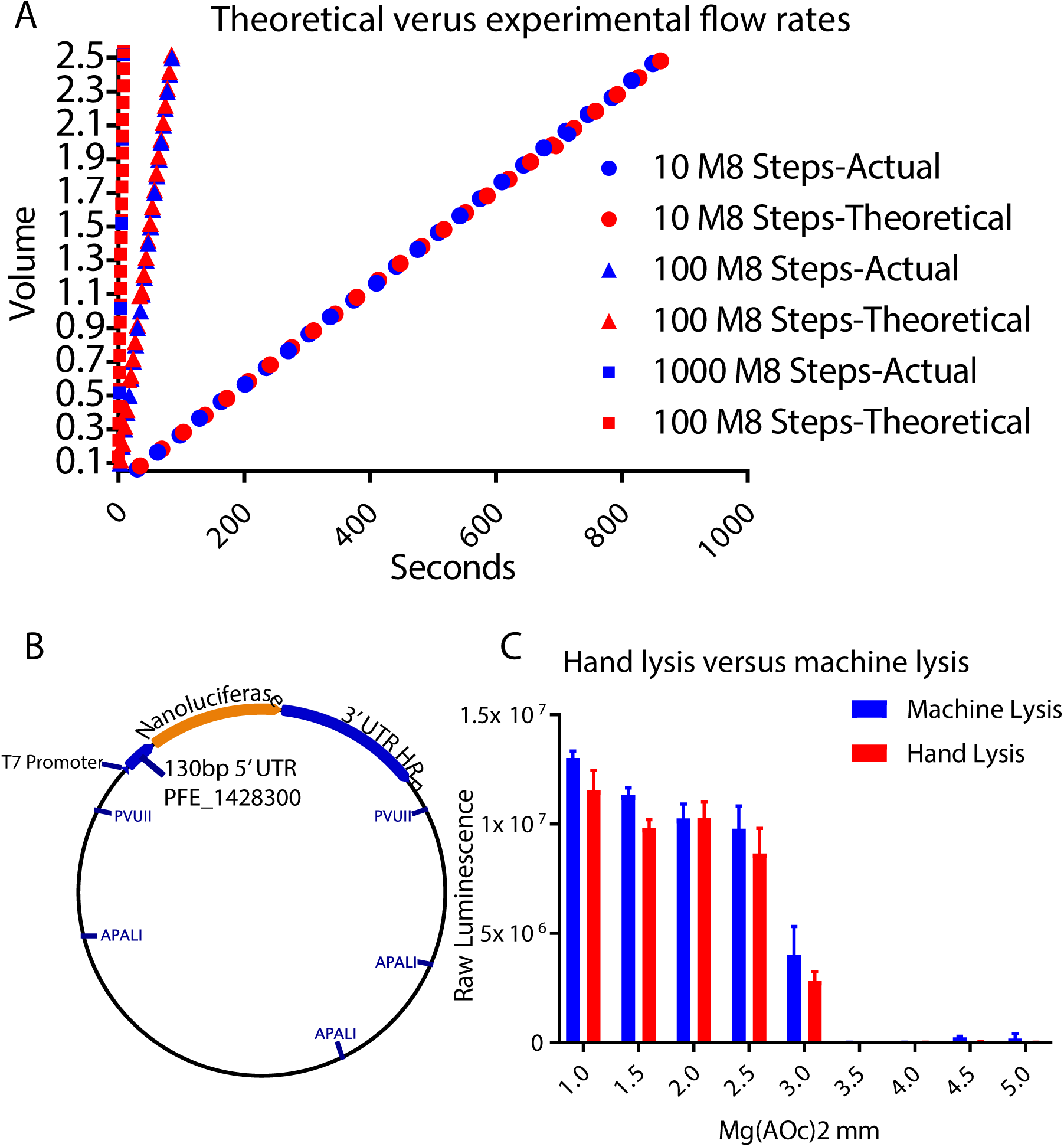
Characterization of the PDSP for lysate generation. A) Dynamic Range of Flow Rate of PDSP with 3mL Syringes. B) Plasmid construct for validation of translational activity in the lysates. C) Translational activity, measured by total luciferase signal, of the lysates generated by hand or the PDSP.

For a T8 threaded rail:

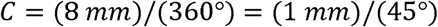

Thus, we can calculate the theoretical flow rate as follows:

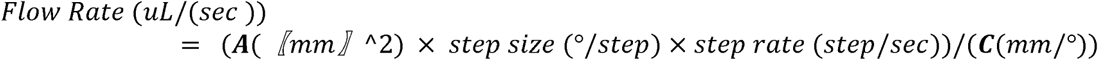

The measured flow rates were consistent with the theoretical flow rates calculated using the manufactured syringe diameters and the stepper motor step sizes. This gives us confidence in the PDSP as an appropriate alternative to commercial syringe pumps. Here we provide the dynamic range of the PDSP in terms of the theoretical minimum and maximum flow rates of various compatible syringe sizes (Table 3).

**Table 3.**
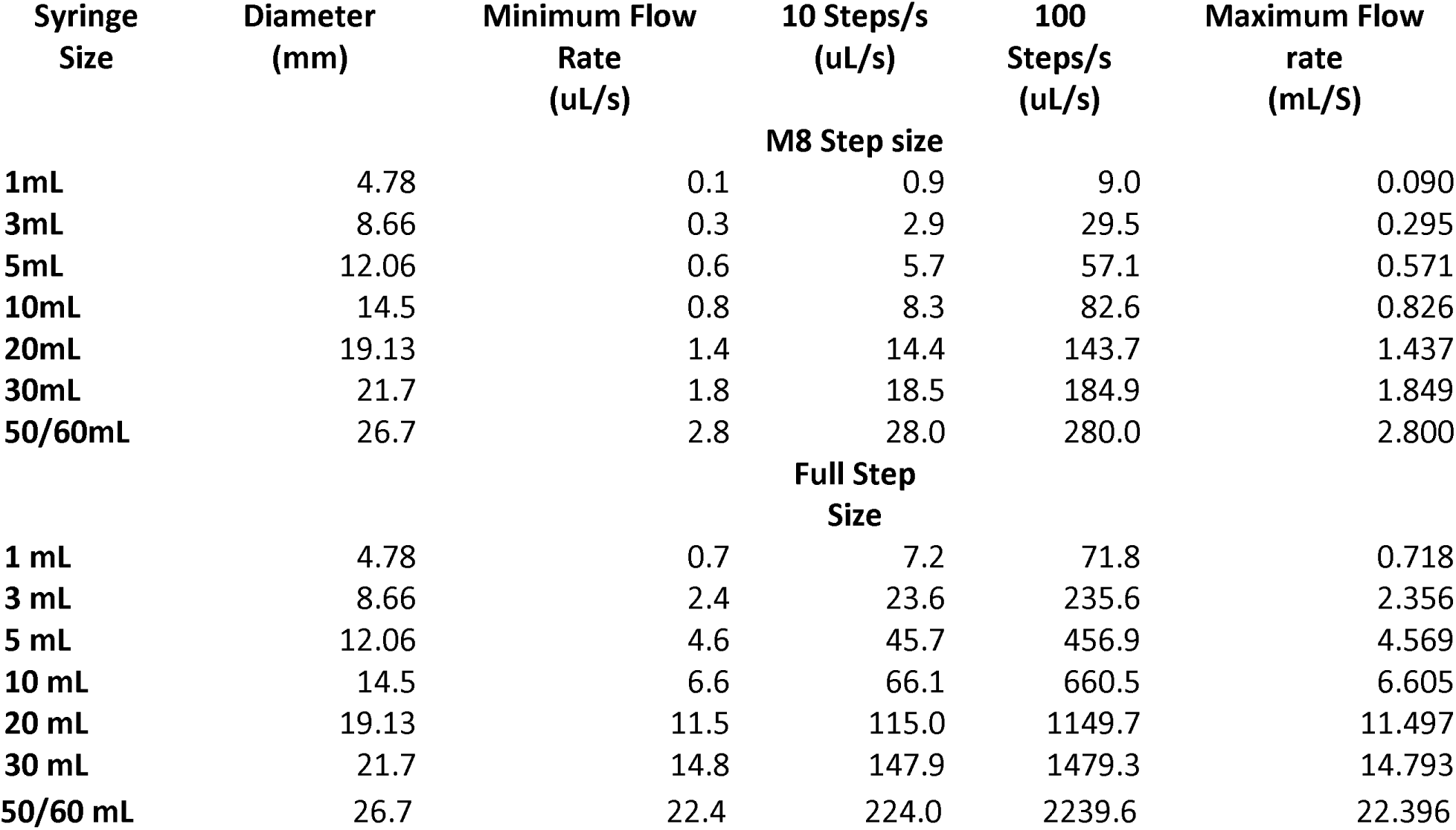
Minimum and Maximum Theoretical Flow Rates by the PDSP Based On Syringe Size.

| Syringe Size | Diameter (mm) | Minimum Flow Rate (uL/s) | 10 Steps/s (uL/s) | 100 Steps/s (uL/s) | Maximum Flow rate (mL/S) |
| --- | --- | --- | --- | --- | --- |
| **M8 Step size** |  |  |  |  |  |
| 1mL | 4.78 | 0.1 | 0.9 | 9.0 | 0.090 |
| 3mL | 8.66 | 0.3 | 2.9 | 29.5 | 0.295 |
| 5mL | 12.06 | 0.6 | 5.7 | 57.1 | 0.571 |
| 10mL | 14.5 | 0.8 | 8.3 | 82.6 | 0.826 |
| 20mL | 19.13 | 1.4 | 14.4 | 143.7 | 1.437 |
| 30mL | 21.7 | 1.8 | 18.5 | 184.9 | 1.849 |
| 50/60mL | 26.7 | 2.8 | 28.0 | 280.0 | 2.800 |
| **Full Step Size** |  |  |  |  |  |
| 1 mL | 4.78 | 0.7 | 7.2 | 71.8 | 0.718 |
| 3 mL | 8.66 | 2.4 | 23.6 | 235.6 | 2.356 |
| 5 mL | 12.06 | 4.6 | 45.7 | 456.9 | 4.569 |
| 10 mL | 14.5 | 6.6 | 66.1 | 660.5 | 6.605 |
| 20 mL | 19.13 | 11.5 | 115.0 | 1149.7 | 11.497 |
| 30 mL | 21.7 | 14.8 | 147.9 | 1479.3 | 14.793 |
| 50/60 mL | 26.7 | 22.4 | 224.0 | 2239.6 | 22.396 |

### 7.2 Validation of the PDSP as a cell homogenizer

To demonstrate the ability of the PDSP to replace manual cell lysis we compared the two methods directly. In brief, *Plasmodium falciparum* parasites were harvested as previously published^1^ and split into two pools, one for manual lysis and one for lysis using the PDSP. We either passed the lysate back-and-forth through the Isobiotec cell homogenizer for 20 cycles by hand or the PDSP controlled by the GUI interface passed the lysate through. Lysates were then collected and centrifuged at 13,000xg for 10 mins at 4C. The supernatant was aliquoted and flash frozen in liquid nitrogen.

One aliquot was taken for each lysis method and used in an *in vitro* translation reaction as previously published with a few modifications^1^. Each 10uL reaction consisted of 7uL of lysate, 10 µM amino acid mixture, 20 mM HEPES/KOH pH 8.0, 75 mM KOAc, a range of 1-5mM Mg(OAc)2, 2 mM DTT, 0.5 mM ATP, 0.1 mM GTP, 20 mM creatine phosphate, 0.2 μg/μl creatine kinase, and 0.5 pmoles of Nanoluciferase reporter RNA. Reactions were done in triplicate, incubated at 37C for 45 mins, and stopped with the addition of 10 uM cycloheximide. The Nanoluciferase reporter RNA consists of the 130 base pairs directly 5’ of PFE_1248300 followed by the Nanoluciferase coding sequence^7^ in the 3’ UTR of HRP. All RNA was generated off of plasmid digested with PVUII and APALl using T7 transcription (Fig. 6B). After transcription, the DNA template was digested with Turbo DNAse (Thermo Fisher) and the RNA was purified using RNA Clean and Concentrate-25 kit (Zymo). Nanoluciferase reactions were performed using Promega’s Nano-Glo Luciferase Assay System. In brief, 10 uL of 1:50 Nanoluciferase substrate:Nanoluciferase buffer was added to each reaction and incubated at room temperature for a minimum of 5 minutes before reading with a 6 second integration on a Promega GloMax-Multi microplate reader.

Our results showed that lysates generated by hand and by the PDSP performed similarly, indicating the PDSP can be used to replace manual lysis for generating *in vitro* translation extracts (Fig. 6C).

### 7.3 Conclusion/Device Overview

We have constructed a programmable syringe pump that can be used for biological life science applications. Not only is the PDSP more affordable than commercially available options, but it is also modular and programmable, allowing the user to customize the device for specific tasks or experiments. Here, we have demonstrated that the PDSP can be used to automate and standardize the time-consuming task of *P. falciparum* lysis. Overall, the PDSP can be used as an affordable and customizable alternative to traditional syringe pumps to automate any number of routine laboratory practices.

## Acknowledgements

We would like to acknowledge Wesley Wu for his contributions to the GUI design and Eric Lam for assisting with the 3D printing.

