## Supplementary material for "Low-Cost Touchscreen Driven Programmable Dual Syringe Pump for Life Science Applications": ExtractMakerGui_current.py

```
#!/usr/bin/python2.7.13

import piplates.MOTORplate as MOTOR
import RPi.GPIO as GPIO
import time
import os
import string
from Tkinter import *
import tkFont
from PIL import Image, ImageTk
import sys


class headerBLOCK:
    def __init__(self,master,r,c):
        self.master=master
        self.tit = Frame(self.master, padx = 4, pady = 4, bd = 2, bg = 'white', relief = 'sunken')
        self.tit.grid(row = r, column = c, columnspan = 2, sticky = N+S+W+E)
         
        ##Create Fonts
        self.title = tkFont.Font(family = 'Helvetica', size = 32, weight = 'bold') 
        self.heading = tkFont.Font(family = 'Helvetica', size = 18, weight = 'bold')           
                
        self.close_button = Button(self.tit, text = 'X', command = self.exitAll).grid(sticky = N+E)  
        
        self.labelt = Label(self.tit, text = 'DeRisi Lab Extract Maker', bg = 'white', fg = '#000666000',
                            padx = 4, pady = 0, width = 32, font = self.title, anchor = 'center').grid(sticky = N+E+S+W) 
        
        self.version_button = Button(self.tit, text = 'Version 1.06', command = self.easterEgg).grid(row=1,column=0, sticky=E+S)

    def easterEgg(self):
        self.window=easterEggWindow(self.master)
        self.master.wait_window(self.window.top)

    def exitAll(self):
        MOTOR.RESET(0)
        GPIO.cleanup(22)
        sys.exit()


class controlGUI:
    global intBITS, intFLAG # Creating varible to ask if the current is changing (binary true or false awnser)
    intBITS = MOTOR.getINTflag0(0)
    intFLAG = 0

    #Sets up visual gui
    def __init__(self,master,r,c):
        self.master=master
        self.initialization=False
        self.sm=Frame(self.master,padx=4,pady=4,bd=2,relief='sunken')
        self.sm.grid(row=r,column=c, sticky=N+S+W+E)
        
        ##Create Fonts
        self.title=tkFont.Font(family='Helvetica', size=25, weight='bold') 
        self.heading=tkFont.Font(family='Helvetica', size=18, weight='bold')
        self.content=tkFont.Font(family='Helvetica', size=12, weight='bold')
        
        
        ##Home Button image
        self.wesHome=Image.open('wes_home.png').convert('RGB')
        self.wesHome = self.wesHome.resize((125,108), Image.ANTIALIAS)
        self.wesHomeDisplay = ImageTk.PhotoImage(self.wesHome)
        #Home button
        self.homeButton=Button(self.sm,image=self.wesHomeDisplay,text='HOME',font=self.heading,compound="center",height=100,width=100,command=self.popupHome)
        self.homeButton.grid(row=0,column=0,rowspan=2,padx=30,pady=4)
        self.homeComplete=0
        
        ##Start/Stop Controls
        self.StartPushSmall=Image.open("StartPushSmall.png").convert("RGB")
        self.StartPushSmall=self.StartPushSmall.resize((125, 108), Image.ANTIALIAS)
        self.StartPushSmallDisplay = ImageTk.PhotoImage(self.StartPushSmall)
        
        self.bigStart=Button(self.sm,image=self.StartPushSmallDisplay,height=108,width=125,command=self.popupStart)
        self.bigStart.image=self.StartPushSmallDisplay
        self.bigStart.grid(row=2,column=0,rowspan=2,padx=30,pady=4)
        
        self.StopPushSmall=Image.open("StopPushSmall.png").convert("RGB")
        self.StopPushSmall=self.StopPushSmall.resize((125, 108), Image.ANTIALIAS)
        self.StopPushSmallDisplay = ImageTk.PhotoImage(self.StopPushSmall)
        
        self.bigStop=Button(self.sm,image=self.StopPushSmallDisplay,height=108,width=125,command=self.stopAll)
        self.bigStop.grid(row=4,column=0,rowspan=2,padx=30,pady=4)
        
        ##Volume Control
        self.labelv=Label(self.sm,text="Volume Control",padx=4,pady=4,font=self.heading,anchor="center")
        self.labelv.grid(row=0,column=1,columnspan=4, sticky=S)
        
        self.vStart=1.5
        self.vLower=Button(self.sm,text="<",fg="black",bg="gray",height=2,width=7,command=self.vdecrease)
        self.vLower.grid(row=1,column=1)
        self.labelvStart=Label(self.sm,text=self.vStart,padx=4,pady=4,font=self.content,anchor="center")
        self.labelvStart.grid(row=1,column=2,columnspan=2,sticky=N+E+S+W)
        self.vUpper=Button(self.sm,text=">",fg="black",bg="gray",height=2,width=7,command=self.vincrease)
        self.vUpper.grid(row=1,column=4)
        
        self.vSet=Button(self.sm,text="Set Total Volume",height=2,width=14,command=self.popupVolume)
        self.vSet.grid(row=2,column=1,columnspan=2,sticky=W)
        self.vReset=Button(self.sm,text="Reset to Default",height=2,width=14,command=self.vdefault)
        self.vReset.grid(row=2,column=3,columnspan=2,sticky=E)
        
        
        ##Cycle Control
        self.labelc=Label(self.sm,text="Cycles Control",padx=4,pady=2,font=self.heading,anchor="center")
        self.labelc.grid(row=3,column=1,columnspan=4,sticky=S)
        
        self.cStart=20
        self.cLower=Button(self.sm,text="<",fg="black",bg="gray",height=2,width=7,command=self.cdecrease)
        self.cLower.grid(row=4,column=1)
        self.labelcStart=Label(self.sm,text=self.cStart,padx=4,pady=4,width=14,font=self.content,anchor="center")
        self.labelcStart.grid(row=4,column=2,columnspan=2,sticky=N+E+W+S)
        self.cUpper=Button(self.sm,text=">",fg="black",bg="gray",height=2,width=7,command=self.cincrease)
        self.cUpper.grid(row=4,column=4,sticky=E)

        self.cSet=Button(self.sm,text="Set Cycle Number",height=2,width=14,command=self.cset)
        self.cSet.grid(row=5,column=1,columnspan=2,sticky=W)
        self.cReset=Button(self.sm,text="Reset to Default",height=2,width=14,command=self.cdefault)
        self.cReset.grid(row=5,column=3,columnspan=2,sticky=E)
        
        
        #Output Information
        self.statusLabel=Label(self.sm,text="Status Overview",padx=4,pady=4,font=self.heading,anchor="center")
        self.statusLabel.grid(row=0,column=5,columnspan=2,sticky=E+S+W)
        
        self.cycleCount=0
        self.cCounterLabel=Label(self.sm,text="Current Cycle",padx=4,pady=4,width=31,font=self.content,anchor="center")
        self.cCounterLabel.grid(row=0,column=5,columnspan=2,sticky=E+S+W)
        self.cycleCountLabel=Label(self.sm,text=self.cycleCount,padx=4,pady=4,width=31,font=self.content,anchor="center")
        self.cycleCountLabel.grid(row=1,column=5,columnspan=2,sticky=N+E+W)
        
        #Time widgets
        self.now="00:00:00"
        self.timeElapsedLabel=Label(self.sm,text="Time Elapsed",padx=4,pady=4,font=self.content,anchor="center")
        self.timeElapsedLabel.grid(row=2,column=5,sticky=S+E+W)
        self.timeUp=Label(self.sm,text=self.now,padx=4,pady=4,font=self.content,anchor="center")
        self.timeUp.grid(row=3,column=5,sticky=N+E+W)
        
        self.remain="00:00:00"
        self.timeRemainLabel=Label(self.sm,text="Time Remaining",padx=4,pady=4,font=self.content,anchor="center")
        self.timeRemainLabel.grid(row=2,column=6,sticky=S+E+W)
        self.timeDown=Label(self.sm,text=self.remain,padx=4,pady=4,font=self.content,anchor="center")
        self.timeDown.grid(row=3,column=6,sticky=N+E+W)

        #Output text widget
        self.text_box=Text(self.sm,padx=4,pady=4,height=6,width=34,state=DISABLED)
        self.text_box.grid(row=4,column=5,columnspan=2,rowspan=2)
        self.scrollb = Scrollbar(self.sm,command=self.text_box.yview)
        self.scrollb.grid(row=4, column=6,rowspan=2,sticky=N+S+E)
        self.text_box['yscrollcommand'] = self.scrollb.set
        
        #Takes standerd output/errors that would go to command line and puts them in white box
        sys.stdout = StdRedirector(self.text_box)
        sys.stderr = StdRedirector(self.text_box)
        
        #Parameters
        self.stat = MOTOR.getSENSORS(0)
        self.homeComplete = 0
        self.vFinal = 0
        self.vCurrent = 0
        self.cFinal = 20
        self.sFinal = 0
        self.direction = 1
        self.dir_dict = {0:'cw', 1:'ccw'}
        
        
        #Job Parameters
        self._job = None
        self._job_vSet = None
        
        
        #Initial information
        print('Please remove all syringes and    press HOME to initialize extract  maker')
        self.disableAllbutHome()
        
    #Change volume functions
    def vset(self):
        global intBITS, intFLAG
        intBITS = MOTOR.getINTflag0(0)
        intFLAG = 0
        
        if self.homeComplete:
            self.bigStart["state"] = "disable"
            self.vFinal=self.vStart
            print self.vFinal-self.vCurrent
            print 'Moving syringe A to', self.vFinal, 'mL'
            
            GPIO.setmode(GPIO.BCM)  
            GPIO.setup(22, GPIO.IN, pull_up_down=GPIO.PUD_UP)  #set up GPIO 22 as an input 
            GPIO.add_event_detect(22, GPIO.FALLING, callback=self.moveDone)  # detect interrupts
            
            if self.vCurrent == 0:
                MOTOR.stepperCONFIG(0,'a','cw','M8',150,0)
                MOTOR.stepperMOVE(0,'a',int(3500*(self.vFinal-self.vCurrent)-1100))
                self.vCurrent=self.vFinal
            elif self.vCurrent > self.vFinal:
                MOTOR.stepperCONFIG(0,'a','ccw','M8',150,0)
                MOTOR.stepperMOVE(0,'a',int(3500*(self.vCurrent-self.vFinal)))
                self.vCurrent=self.vFinal
            elif self.vCurrent < self.vFinal:
                MOTOR.stepperCONFIG(0,'a','cw','M8',150,0)
                MOTOR.stepperMOVE(0,'a',int(3500*(self.vFinal-self.vCurrent)))
                self.vCurrent=self.vFinal
            root.after(100,self.checkvSet)
        else:
            print "Please home motors first"
    
    def checkvSet(self):
        global intBITS, intFLAG
        #print intBITS
        if intFLAG:
            intBITS = MOTOR.getINTflag0(0)
            intFLAG = 0
            if not intBITS&0x10:
            #if intBITS&0x20:
                self.sync_stop()
                print 'Syringe set at', self.vCurrent, 'mL'
                self.bigStart["state"] = "normal"
                self.cancel()
                GPIO.cleanup(22)
        else:
            self._job=root.after(100,self.checkvSet)

    
    def popupVolume(self):
        self.window=PopupWindowVolume(self.master)
        self.master.wait_window(self.window.top)
        root.after(1,self.vset)
    
    #This sets the default button values
    def vdefault(self):
        if self.homeComplete:
            #Set volume counter back to default value of 1.5mL
            #This does not change the current syringe position
            self.vStart=1.5
            self.labelvStart.config(text=self.vStart)
        else:
            print "Please home motors first"
    
    def vdecrease(self):
        #Decrease volume by 0.1mL to a minimum of 1.0mL
        if self.homeComplete:
            if self.vStart > 0.5:
                self.vStart-=0.1
                self.vStart = round(self.vStart,1)
                self.labelvStart.config(text=self.vStart)
                #print(self.vStart)
        else:
            print "Please home motors first"
            
    def vincrease(self):
        #Increase the volumeby 0.1mL to a maximum of 3.0mL
        if self.homeComplete:
            if self.vStart < 3:
                self.vStart+=0.1
                self.vStart = round(self.vStart,1)
                self.labelvStart.config(text=self.vStart)
                #print(self.vStart)
        else:
            print "Please home motors first"
    
    #Change cycles functions
    def cset(self):
        if self.homeComplete:
            self.cFinal=self.cStart
            print "Set cycle number to",self.cFinal
        else:
            print "Please home motors first"
        
    def cdefault(self):
        if self.homeComplete:
            #Reset cycle number to a default value of 20
            self.cStart=20
            self.labelcStart.config(text=self.cStart)  
        else:
            print "Please home motors first"
            
    def cdecrease(self):
        if self.homeComplete:
            #Decrease cycle number by 1
            if self.cStart > 1:
                self.cStart-=1
                self.labelcStart.config(text=self.cStart)
                #print(self.vStart)
        else:
            print "Please home motors first"
    
    def cincrease(self):
        if self.homeComplete:            
            #Increase cycle number by 1
            self.cStart+=1
            self.labelcStart.config(text=self.cStart)
            #print(self.vStart)
        else:
            print "Please home motors first"
            
            
    #Home functions
    def jogHome(self):
        #Initialize homing variables
        self.motorA = 1
        self.motorB = 1
        self.homeButton["state"] = "disabled" 
        #Configure motor parameters for jogging
        MOTOR.stepperCONFIG(0,'a','ccw','M8',150,0)
        MOTOR.stepperCONFIG(0,'b','ccw','M8',150,0)
        #Move steppers A and B towards switches
        MOTOR.stepperJOG(0,'a')
        MOTOR.stepperJOG(0,'b')
        #Wait for limit switch signal
        root.after(1,self.findHome)

    def findHome(self):
        #Sample every 100ms
        self._job = root.after(100,self.findHome)
        self.stat = MOTOR.getSENSORS(0)
        #If limit switch A is hit, stop and reverse
        if not (self.stat&0x01):
            MOTOR.stepperSTOP(0,'a')
            MOTOR.stepperCONFIG(0,'a','cw','M8',100,0)
            MOTOR.stepperMOVE(0,'a',150)
            MOTOR.stepperOFF(0,'a')
            self.motorA = 0
            print "MOTOR B homing..."
        #If limit switch B is hit, stop and reverse
        if not (self.stat&0x04):
            MOTOR.stepperSTOP(0,'b')
            MOTOR.stepperCONFIG(0,'b','cw','M8',100,0)
            MOTOR.stepperMOVE(0,'b',150)
            MOTOR.stepperOFF(0,'b')
            self.motorB = 0
            print "MOTOR A homing..."
        #When both motors are stopped, exit after looping
        if (self.motorA == 0 and self.motorB == 0):
            self.enableAll()
            self.vCurrent = 0
            self.cycleCount = 0
            self.homeComplete = 1
            print"Home complete"
            self.homeButton["state"] = "normal"
            root.after_cancel(self._job)
            #self.cancel()
        
      
    def popupHome(self):
        self.window=PopupWindowHome(self.master)
        self.master.wait_window(self.window.top)
        root.after(1,self.jogHome)
      
    #Lysate making functions
    def popupStart(self):
        self.window=PopupWindowStart(self.master)
        self.master.wait_window(self.window.top)
        root.after(1,self.makeLysate)

    def makeLysate(self):
        global intBITS, intFLAG
        if self.homeComplete:
            if self.vFinal == 0:
                print('Set Lysate Volume')
            if self.cFinal == 0:
                print('Set Cycle Number')
            else:
                GPIO.setmode(GPIO.BCM)  
                GPIO.setup(22, GPIO.IN, pull_up_down=GPIO.PUD_UP)  #set up GPIO 22 as an input 
                GPIO.add_event_detect(22, GPIO.FALLING, callback=self.moveDone)  # detect interrupts
                self.disableAllbutHome()
                self.homeButton["state"] = "disabled" 
                print 'Making', self.vFinal, 'mL of lysate in', self.cFinal, 'cycles'
                self.sync_config(self.direction,120,0)
                self.start = time.time()
                time.clock()
                self.remainSeconds = int(self.cFinal*((3500*self.vFinal)-1400)/60)
                root.after(1,self.oneCycle)
        else:
            print('Remove syringes and home motors')
    
    def oneCycle(self):
        global intBITS, intFLAG
        root.after(100,self.limitSwitch)
        if self.cycleCount < self.cFinal:
            self.sync_move(int((3500*self.vFinal)-1400))
            #print "moving", str((3500*self.vFinal)-1400)
            self.cycleCountLabel.config(text=self.cycleCount)
            root.after(1000, self.update_clock)
            root.after(1000, self.update_timer)
            self._job = root.after(100,self.doCycle)
        elif self.cycleCount == self.cFinal:
            if self.cycleCount == 0:
                GPIO.cleanup(22)
                print('Process aborted. Check last parameters')
            else:
                GPIO.cleanup(22)
                print 'Completed Lysing of', self.vFinal, 'mL of lysate in', self.cFinal, 'cycles'
                self.cycleCountLabel.config(text=self.cycleCount)
                self.stopAll()
                root.after(1000, self.update_clock_end)
        else:
            GPIO.cleanup(22)
            print('Not sure what happened, but RESET!')
            self.stopAll
                 
    def doCycle(self):

        global intBITS, intFLAG
        if intFLAG:
            intFLAG = 0
            intBITS = MOTOR.getINTflag0(0)
            if not intBITS&0x20:
                print "stopping"
                self.sync_stop()
                self.direction = self.toggle_direction(self.direction)
                self.sync_config(self.direction,120,0)
                print "changing directions"
                self.cycleCount += 0.5
                print self.cycleCount
                
        self._job=root.after(100,self.oneCycle)
    
    def toggle_direction(self,dir_val):
        return (1-dir_val)
    
    def sync_config(self,dir,rate,acc):
        MOTOR.stepperCONFIG(0,'a',self.dir_dict[dir],'M8',rate,acc)
        MOTOR.stepperCONFIG(0,'b',self.dir_dict[self.toggle_direction(dir)],'M8',rate,acc)
    
    def sync_move(self,steps):
        MOTOR.stepperMOVE(0,'a',steps)
        MOTOR.stepperMOVE(0,'b',steps)
    
    def sync_stop(self):
        MOTOR.stepperSTOP(0,'a')
        MOTOR.stepperSTOP(0,'b')
        
    #Interrupt functions
    def moveDone(self, channel):
        # signal to main program that an int has occurred
        global intBITS, intFLAG
        intBITS = MOTOR.getINTflag0(0)
        intFLAG = 1
        
    def limitSwitch(self):
        self.stat = MOTOR.getSENSORS(0)
        if not (self.stat&0x01) or not (self.stat&0x04):
            MOTOR.RESET(0)
            print('Limit switches hit, stopping')
            root.after_cancel(self._job)
            #self.cancel()
            GPIO.cleanup(22)
            self.stopAll()
    
    #Stopping functions
    def stopAll(self):
        #global intBITS, intFLAG
        #intBITS = MOTOR.getINTflag0(0)
        #intFLAG = 0
        self.cancel()
        #root.after_cancel(self._job)
        MOTOR.RESET(0)
        GPIO.cleanup(22)
        self.homeComplete = 0
        self.vFinal = 0
        self.vCurrent = 0
        self.cFinal = 20
        self.cCurrent = 0
        self.cycleCount = 0
        self.direction = 1
        self.disableAllbutHome()
        self.homeButton["state"] = "normal"
        self.resetClocks()
        print('Stopping all processes')
        print('Please re-Home and reset volume and cycles')
        
    def cancel(self):
        if self._job is not None:
            root.after_cancel(self._job)
            self._job = None
        if self._job_vSet is not None:
            root.after_cancel(self._job_vSet)
            self.job_vSet = None

    
    def disableAllbutHome(self):
        self.bigStart["state"] = "disabled" 
        #self.bigStop["state"] = "disabled"
        
        self.vLower["state"] = "disabled" 
        self.vUpper["state"] = "disabled" 
        self.vSet["state"] = "disabled"
        self.vReset["state"] = "disabled"
        
        self.cLower["state"] = "disabled" 
        self.cUpper["state"] = "disabled" 
        self.cSet["state"] = "disabled" 
        self.cReset["state"] = "disabled"
        
    def enableAll(self):
        #self.bigStart["state"] = "normal" 
        #self.bigStop["state"] = "normal"
        
        self.vLower["state"] = "normal" 
        self.vUpper["state"] = "normal" 
        self.vSet["state"] = "normal"
        self.vReset["state"] = "normal"
        
        self.cLower["state"] = "normal" 
        self.cUpper["state"] = "normal" 
        self.cSet["state"] = "normal" 
        self.cReset["state"] = "normal"

    #Clock functions
    def update_clock(self):
        self.now = int(time.time() - self.start)
        self.now = self.getTime(self.now)
        self.timeUp.configure(text=self.now)
        
    def update_clock_end(self):
        self.now = int(time.time() - self.start)
        self.now = self.getTime(self.now)
        self.timeUp.configure(text=self.now)
        root.after(1000,self.update_clock_end)

    def getTime(self,seconds):
        self.m, self.s = divmod(seconds, 60)
        self.h, self.m = divmod(self.m, 60)
        return "%d:%02d:%02d" % (self.h, self.m, self.s)
    
    def update_timer(self):
        if self.remain <= 0:
            self.timeDown.configure(text="time's up!")
        else:
            self.remain = int(self.remainSeconds - (time.time() - self.start))
            self.remain = self.getTime(self.remain)
            self.timeDown.configure(text=self.remain)
   
    def resetClocks(self):
        self.now="00:00:00"
        self.timeUp.configure(text=self.now)
        self.remain="00:00:00"
        self.timeDown.configure(text=self.remain)
   
class StdRedirector():
    def __init__(self, text_widget):
        self.text_space = text_widget

    def write(self, string):
        self.text_space.config(state=NORMAL)
        self.text_space.insert("end", string)
        self.text_space.see("end")
        self.text_space.config(state=DISABLED)

class PopupWindowHome(object):
    def __init__(self,master):
        top=self.top=Toplevel(master)
        self.l=Label(top,text='Please remove all syringes from the lysate maker',font=tkFont.Font(family='Helvetica', size=24),padx=4,pady=4)
        self.l.pack()
        self.b=Button(top,text='All Clear!',height=4,width=12,padx=4,pady=4,font=tkFont.Font(family='Helvetica', size=12),command=self.cleanup)
        self.b.pack()
    def cleanup(self):
        self.top.destroy()

class PopupWindowStart(object):
    def __init__(self,master):
        top=self.top=Toplevel(master)
        self.l=Label(top,text='Secure your syringe/lysate maker in place',font=tkFont.Font(family='Helvetica', size=24),padx=4,pady=4)
        self.l.pack()
        self.b=Button(top,text='Ready to Lyse!',height=4,width=12,padx=4,pady=4,font=tkFont.Font(family='Helvetica', size=12),command=self.cleanup)
        self.b.pack()
    def cleanup(self):
        self.top.destroy()

class PopupWindowVolume(object):
    def __init__(self,master):
        top=self.top=Toplevel(master)
        self.l=Label(top,text='Remove syringe for motors to move to set volume',font=tkFont.Font(family='Helvetica', size=24),padx=4,pady=4)
        self.l.pack()
        self.b=Button(top,text='Move Motors!',height=4,width=12,padx=4,pady=4,font=tkFont.Font(family='Helvetica', size=12),command=self.cleanup)
        self.b.pack()
    def cleanup(self):
        self.top.destroy()

class easterEggWindow(object):
    def __init__(self,master):
        top=self.top=Toplevel(master)
        ##Image Import
        self.easteregg=Image.open('easteregg.jpg').convert('RGB')
        self.easteregg=self.easteregg.resize((403,302), Image.ANTIALIAS)
        self.easterDisplay = ImageTk.PhotoImage(self.easteregg)
        self.l=Label(top,image=self.easterDisplay)
        self.l.pack()
        self.b=Button(top,text='You can do it!!', height=2, width=12, padx=4,pady=4,font=tkFont.Font(family='Helvetica', size=12),command=self.cleanup)
        self.b.pack()
    def cleanup(self):
        self.top.destroy()

#Reset motors
MOTOR.RESET(0)  
MOTOR.enablestepSTOPint(0,'a')
MOTOR.enablestepSTOPint(0,'b')
MOTOR.intEnable(0)

#Initialize Tkinter GUI
root = Tk()
root.config(bg="black")
root.attributes("-fullscreen", TRUE)
swidth=root.winfo_screenwidth()
sheight=root.winfo_screenheight()
#print tkFont.families()
container=Frame(root,bg="white")
container.place(relx=0.5, rely=0.5, anchor=CENTER)

title = headerBLOCK(container,0,0)
control = controlGUI(container,1,0)

root.mainloop()
```
