## Supplementary material for "Low-Cost Touchscreen Driven Programmable Dual Syringe Pump for Life Science Applications": GUI_pygubu.py

```
import piplates.MOTORplate as MOTOR
import RPi.GPIO as GPIO
import time
import os
import string
import Tkinter as tk
import tkFont
import pygubu
import sys


class Application:
    def __init__(self, master):

        #1: Create a builder
        self.builder = builder = pygubu.Builder()

        #2: Load an ui file
        builder.add_from_file('guiDesign_pygubu.ui')

        #3: Create the widget using a master as parent
        self.mainwindow = builder.get_object('containerFrame', master)

        #4 Connect method callbacks
        builder.connect_callbacks(self)
        
        self.cStart=5
        self.vStart=2
        
    def exitWindow(self):
        sys.exit()
        
    def vdecrease(self):
        if self.vStart > 0.5:
            self.vStart-=0.1
            self.vStart = round(self.vStart,1)
            self.vCurrent.config(text=self.vStart)
            #print(self.vStart)
        
    def vincrease(self):
        if self.vStart < 3:
            self.vStart+=0.1
            self.vStart = round(self.vStart,1)
            self.vCurrent.config(text=self.vStart)
            #print(self.vStart)
        

    def cdecrease(self):
        if self.cStart > 1:
            self.cStart-=1
            self.cCurrent.config(text=self.cStart)
            #print(self.vStart)
        
    def cincrease(self):
        self.cStart+=1
        print(self.cStart)
        self.cCurrent.config(text=self.cStart)
        #print(self.vStart)
    
    def makeLysate(self):
        self.homeComplete=1
        print("making lysate")
    
    def stopAll(self):
        MOTOR.RESET(0)
        self.homeComplete=0
        print("stopping")

        

if __name__ == '__main__':
    root = tk.Tk()
    app = Application(root)
    swidth=root.winfo_screenwidth()
    sheight=root.winfo_screenheight()
    root.config(bg="black")

    
    
    root.mainloop()   
```
