## Supplementary material for "Low-Cost Touchscreen Driven Programmable Dual Syringe Pump for Life Science Applications": Home_JL.py

```
import piplates.MOTORplate as MOTOR
import RPi.GPIO as GPIO
import time
import os
import string
from Tkinter import *
import tkFont

print MOTOR.getID(0)

#flush out any old interrupts
#dummy = MOTOR.getINTflag0(1)
#intFLAG=0
#intITS=0

#def moveDone(channel):
#    global intBITS, intFLAG
#    intBITS=MOTOR.getINTflag0(0)
#    intFLAG=1

MOTOR.RESET(0)

#configure steppers A and B for homing function
MOTOR.stepperCONFIG(0,'a','ccw','M8',100,0)
MOTOR.stepperCONFIG(0,'b','ccw','M8',100,0)


#move steppers A and B towards switches
MOTOR.stepperJOG(0,'a')
MOTOR.stepperJOG(0,'b')

#GPIO.setmode(GPIO.BCM)
#GPIO.setup(22, GPIO.IN, pull_up_down=GPIO.PUD_UP)
#GPIO.add_event_detect(22, GPIO.FALLING, callback=moveDone)
#allow interrupts when a motor stops
#MOTOR.enablestepSTOPint(0,'a')
#MOTOR.enablestepSTOPint(0,'b')
#MOTOR.intEnable(0)


flag=1
motorA=1
motorB=1


while(flag):
    time.sleep(0.1)
    stat=MOTOR.getSENSORS(0)
    if not (stat&0x01):
        MOTOR.stepperSTOP(0,'a')
        time.sleep(0.1)
        MOTOR.stepperCONFIG(0,'a','cw','M8',100,0)
        MOTOR.stepperMOVE(0,'a',50)
        MOTOR.stepperOFF(0,'a')
        motorA=0
        print "MOTOR A stopped"
    if not (stat&0x04):
        MOTOR.stepperSTOP(0,'b')
        MOTOR.stepperCONFIG(0,'b','cw','M8',100,0)
        MOTOR.stepperMOVE(0,'b',50)
        MOTOR.stepperOFF(0,'b')
        motorB=0
        print "MOTOR B stopped"
    if (motorA==0 and motorB==0):
        flag = 0
        print"End Stop Reached"
        
```
