## Supplementary material for "Low-Cost Touchscreen Driven Programmable Dual Syringe Pump for Life Science Applications": MOTORdemoOOP.py

```
import piplates.MOTORplate as MOTOR 
import time
import os
import string
from Tkinter import *
import tkFont

class titleBLOCK:
    def __init__(self,master,r,c):
        self.master=master
        self.tit=Frame(self.master,padx=4,pady=4,bd=2,bg='white',relief='sunken')
        self.tit.grid(row=r,column=c, columnspan=2,sticky=N+S+W+E)    
         
        ##Create Fonts
        self.title = tkFont.Font(family='Helvetica', size=40, weight='bold') 
        self.heading = tkFont.Font(family='Helvetica', size=18, weight='bold')       

        self.logo = PhotoImage(file = '3D-ppLogo.gif')
        self.pp=Label(self.tit,image=self.logo,anchor="center").grid() 
        
        self.labelt = Label(self.tit, text="MOTORplate Controller", bg='white',fg='#000666000',padx=4,pady=4,font=self.title,anchor="center")
        self.labelt.grid(sticky=W+E)      
        

        #self.pp.bind("<ButtonRelease-1>", self.shutdown)
        
        self.close_button = Button(self.tit, text="X", command=root.quit).grid(sticky=E)  
        
        def shutdown(self,event):
            print "clicked at", event.x, event.y
        
        
class stepperGUI:
    def __init__(self,master,title,stepper,r,c):
        self.stepState=0       #stopped
        smColor="yellow"
        slwidth=30
        slength=120
        self.mVal=stepper
        self.master=master
        self.sm=Frame(self.master,padx=4,pady=4,bd=2,relief='sunken')
        self.sm.grid(row=r,column=c, rowspan=3, sticky=N+S+W+E)
        
        ##Create Fonts
        self.title = tkFont.Font(family='Helvetica', size=25, weight='bold') 
        self.heading = tkFont.Font(family='Helvetica', size=18, weight='bold')
        
        ##Title
        self.labelt = Label(self.sm, text=title, padx=4,pady=4, bg=smColor, fg='black',font=self.title)
        self.labelt.grid(row=0,column=0,columnspan=2,sticky=W+E)
        
        ##Step Rate Control
        self.labels = Label(self.sm, text="Step Rate",padx=4,pady=4,font=self.heading)
        self.labels.grid(row=1,column=0)
        self.rate=IntVar()
        self.rate.set(0)
        self.RateSet=Scale(self.sm,variable=self.rate,from_=2000,to=0,width=slwidth,length=slength)
        self.RateSet.bind("<ButtonRelease-1>", self.ratedelta)
        self.RateSet.grid(row=2,column=0,rowspan=2)
 
        ##Direction Control
        self.labeld = Label(self.sm, text="Direction",padx=4,pady=4,font=self.heading)
        self.labeld.grid(row=1,column=1,sticky=W)       
        self.direction = IntVar()
        self.direction.set(0)
        self.cwb = Radiobutton(self.sm, text='Clockwise', variable=self.direction, value=0)
        self.ccwb = Radiobutton(self.sm, text=' Counter Clockwise', variable=self.direction, value=1)
        self.direction.set(0)
        self.cwb.grid(row=2,column=1,sticky=W)
        self.ccwb.grid(row=3,column=1,sticky=W)

        ##Acceleration Control
        self.labela = Label(self.sm, text="Acceleration",padx=4,pady=4,font=self.heading)
        self.labela.grid(row=4,column=0)
        self.acc=DoubleVar()
        self.acc.set(0.0)
        self.accSet=Scale(self.sm,variable=self.acc,from_=5.0,to=0.0, resolution=0.1,width=slwidth,length=slength)
        self.accSet.grid(row=5,column=0,rowspan=4)

        ##Step Size Control
        rStart = 4     
        self.labeld = Label(self.sm, text="Step Size",padx=4,pady=4,font=self.heading)
        self.labeld.grid(row=rStart,column=1,sticky=W) 
        rStart+=1
        self.ss = IntVar()
        self.ss.set(2)  # initializing the choice: Full
        sizes = [
            ("Full",0),
            ("Half",1),
            ("1/4 - microstep",2),
            ("1/8 - microstep",3)
            ] 
        
        for txt, val in sizes:
            Radiobutton(self.sm, 
                text=txt, 
                variable=self.ss, 
                value=val).grid(row=rStart+val,column=1,sticky=W)

        ##Step Count Control
        self.labelstep = Label(self.sm, text="Step Count",padx=4,pady=4,font=self.heading)
        self.labelstep.grid(row=9,column=0)
        self.steps=IntVar()
        self.steps.set(0)
        self.StepSet=Scale(self.sm,variable=self.steps,from_=2000,to=0,width=slwidth,length=slength)
        self.StepSet.grid(row=10,column=0,rowspan=3)                

        ##Jog Button
        self.startButton=Button(self.sm,text="JOG",fg="black",bg="green",height=3, width=6,command=self.jog)
        self.startButton.grid(row=10,column=1)
 
        ##Move Button
        self.startButton=Button(self.sm,text="MOVE",fg="white",bg="blue",height=3, width=6,command=self.move)
        self.startButton.grid(row=11,column=1) 
        
        ##Stop Button
        self.stopButton=Button(self.sm,text="STOP",fg="white",bg="red",height=3, width=6,command=self.stop)
        self.stopButton.grid(row=12,column=1)  

        ##Off Button
        self.stopButton=Button(self.sm,text="OFF",fg="black",bg="yellow",height=3, width=6,command=self.off)
        self.stopButton.grid(row=13,column=1) 
        
        self.stop()   #ensure motor is off at start
        self.off()   #ensure motor is off at start
 
        
    def ratedelta(self,val):
        if (self.stepState==1):
            MOTOR.stepperRATE(0,self.mVal,self.rate.get())

    def jog(self):
        if (self.direction.get() == 0):
            dir='cw'
        else:
            dir='ccw'
        MOTOR.stepperCONFIG( 0,self.mVal,dir,self.ss.get(),self.rate.get(),self.acc.get())
        MOTOR.stepperJOG(0,self.mVal)
        self.stepState=1
        
    def move(self):
        if (self.direction.get() == 0):
            dir='cw'
        else:
            dir='ccw'
        MOTOR.stepperCONFIG(0,self.mVal,dir,self.ss.get(),self.rate.get(),self.acc.get())
        MOTOR.stepperMOVE(0,self.mVal,self.steps.get())
        self.stepState=2
        
    def stop(self):
        MOTOR.stepperSTOP(0,self.mVal)    
        self.stepState=0            

    def off(self):
        MOTOR.stepperOFF(0,self.mVal)    
        self.stepState=0         

class dcGUI:   
    def __init__(self,master,title,mN,r,c):
        self.dcState=0       #stopped
        dcColor="red"
        slwidth=30
        self.mNum=mN
        self.master=master
        self.dcm=Frame(master,padx=4,pady=4,bd=2,relief='sunken')
        self.dcm.grid(row=r,column=c, columnspan=1, rowspan=2, sticky=N+S+W+E)
        
        ##Font Setup
        self.title = tkFont.Font(family='Helvetica', size=25, weight='bold')       
        self.heading = tkFont.Font(family='Helvetica', size=18, weight='bold') 
        
        ##Title
        self.labelt = Label(self.dcm, text=title,  padx=4,pady=4, fg="white", bg=dcColor, font=self.title)
        self.labelt.grid(row=0,column=0,columnspan=2,sticky=W+E)
        
        #Speed Control
        self.labels = Label(self.dcm, text="Speed",padx=4,pady=4,font=self.heading)
        self.labels.grid(row=1,column=0)
        self.speed=IntVar()
        self.speed.set(0)
        self.SpeedSet=Scale(self.dcm,variable=self.speed,from_=100,to=0,width=slwidth)
        self.SpeedSet.bind("<ButtonRelease-1>", self.speeddelta)
        self.SpeedSet.grid(row=2,column=0,rowspan=2)
        
        ##Direction Control
        self.labeld = Label(self.dcm, text="Direction",padx=4,pady=4,font=self.heading)
        self.labeld.grid(row=1,column=1,sticky=W)       
        self.direction = IntVar()
        self.direction.set(0)
        self.cwb = Radiobutton(self.dcm, text='Clockwise', variable=self.direction, value=0)
        self.ccwb = Radiobutton(self.dcm, text=' Counter Clockwise', variable=self.direction, value=1)
        self.direction.set(0)
        self.cwb.grid(row=2,column=1,sticky=W)
        self.ccwb.grid(row=3,column=1,sticky=W)
        
        ##Acceleration Control
        self.labela = Label(self.dcm, text="Acceleration",padx=4,pady=4,font=self.heading)
        self.labela.grid(row=4,column=0)
        self.acc=DoubleVar()
        self.acc.set(0.0)
        self.accSet=Scale(self.dcm,variable=self.acc,from_=5.0,to=0.0, resolution=0.1,length=100,width=slwidth)
        self.accSet.grid(row=5,column=0,rowspan=2)
        
        ##RPM Display
        self.rpm=StringVar()
        self.rpm.set('0')        
        self.labelt = Label(self.dcm, text="RPM",padx=4,pady=4,font=self.heading)
        self.labelt.grid(row=7,column=0,columnspan=2) 
        self.labelt = Label(self.dcm, textvariable=self.rpm ,padx=4,pady=4,font=self.heading)
        self.labelt.grid(row=8,column=0,columnspan=2)        
        
        ##Start Button
        self.startButton=Button(self.dcm,text="GO",fg="black",bg="green",height=3, width=6,command=self.go)
        self.startButton.grid(row=5,column=1)
        
        ##Stop Button
        self.stopButton=Button(self.dcm,text="STOP",fg="white",bg="red",height=3, width=6,command=self.stop)
        self.stopButton.grid(row=6,column=1)
        
        self.UpdateRPM() 
        
    def go(self):
        if (self.direction.get() == 0):
            dir='cw'
        else:
            dir='ccw'
        MOTOR.dcCONFIG(1,self.mNum,dir,self.speed.get(),self.acc.get())
        MOTOR.dcSTART(1,self.mNum)
        self.dcState=1
        
    def stop(self):
        MOTOR.dcSTOP(1,self.mNum)    
        self.dcState=0
    
    def speeddelta(self,val):
        #print self.dcState,self.speed.get()
        if (self.dcState):
            MOTOR.dcSPEED(1,self.mNum,self.speed.get())
        

    def UpdateRPM(self):
        t=MOTOR.getTACHfine(1,self.mNum)
        self.rpm.set(str(t*60/300))
        self.dcm.after(1000,self.UpdateRPM)    

    

MOTOR.RESET(0)
MOTOR.RESET(1)    
        
root = Tk()
root.config(bg="black")
root.attributes("-fullscreen", True)
swidth=root.winfo_screenwidth()
sheight=root.winfo_screenheight()
#print tkFont.families()
container=Frame(root,bg="white")
container.place(relx=0.5, rely=0.5, anchor=CENTER)


ma_gui = stepperGUI(container,"Stepper Motor A",'a',1,1)
mb_gui = stepperGUI(container,"Stepper Motor B",'b',1,2)
m1_gui = dcGUI(container,"DC Motor 1",1,0,0)
m2_gui = dcGUI(container,"DC Motor 2",2,2,0)
m3_gui = dcGUI(container,"DC Motor 3",3,0,3)
m4_gui = dcGUI(container,"DC Motor 4",4,2,3)
title = titleBLOCK(container,0,1)


root.mainloop()
```
