## Supplementary material for "Low-Cost Touchscreen Driven Programmable Dual Syringe Pump for Life Science Applications": New_GUI_JL2.py

```
import piplates.MOTORplate as MOTOR
import RPi.GPIO as GPIO
import time
import os
import string
from Tkinter import *
import tkFont
from PIL import Image, ImageTk
import sys


class headerBLOCK:
    def __init__(self,master,r,c):
        self.master=master
        self.tit=Frame(self.master,padx=4,pady=4,bd=2,bg='white',relief='sunken')
        self.tit.grid(row=r,column=c, columnspan=2,sticky=N+S+W+E)
         
        ##Create Fonts
        self.title = tkFont.Font(family='Helvetica', size=32, weight='bold') 
        self.heading = tkFont.Font(family='Helvetica', size=18, weight='bold')           
                
        self.close_button = Button(self.tit, text="X", command=root.quit).grid(sticky=N+E)  
        
        self.labelt = Label(self.tit, text="DeRisi Lab Extract Maker", bg='white',fg='#000666000',padx=4,pady=4,width=32,font=self.title,anchor="center")
        self.labelt.grid(sticky=N+E+S+W) 

        
        def shutdown(self,event):
            print "clicked at", event.x, event.y


class controlGUI:
    global intBITS, intFLAG
    intBITS=MOTOR.getINTflag0(0)
    intFLAG=0

            
    def __init__(self,master,r,c):
        self.master=master
        self.initialization=False
        self.sm=Frame(self.master,padx=4,pady=4,bd=2,relief='sunken')
        self.sm.grid(row=r,column=c, sticky=N+S+W+E)
        
        ##Create Fonts
        self.title=tkFont.Font(family='Helvetica', size=25, weight='bold') 
        self.heading=tkFont.Font(family='Helvetica', size=18, weight='bold')
        self.content=tkFont.Font(family='Helvetica', size=12, weight='bold')
        
        
        ##Home Button
        self.homeButton=Button(self.sm,text="HOME",fg="black",bg="blue",height=6,width=16,command=self.jogHome)
        self.homeButton.grid(row=0,column=0,rowspan=2,padx=30,pady=4)
        self.homeComplete=0
        
        ##Start/Stop Controls
        self.StartPushSmall=Image.open("StartPushSmall.png").convert("RGB")
        self.StartPushSmall=self.StartPushSmall.resize((125, 108), Image.ANTIALIAS)
        self.StartPushSmallDisplay = ImageTk.PhotoImage(self.StartPushSmall)
        
        self.bigStart=Button(self.sm,image=self.StartPushSmallDisplay,height=108,width=125,command=self.makeLysate)
        self.bigStart.image=self.StartPushSmallDisplay
        self.bigStart.grid(row=2,column=0,rowspan=2,padx=30,pady=4)
        #self.bigStart.pack
        
        self.StopPushSmall=Image.open("StopPushSmall.png").convert("RGB")
        self.StopPushSmall=self.StopPushSmall.resize((125, 108), Image.ANTIALIAS)
        self.StopPushSmallDisplay = ImageTk.PhotoImage(self.StopPushSmall)
        
        self.bigStop=Button(self.sm,image=self.StopPushSmallDisplay,height=108,width=125,command=self.stopAll)
        self.bigStop.grid(row=4,column=0,rowspan=2,padx=30,pady=4)
        
        ##Volume Control
        self.labelv=Label(self.sm,text="Volume Control",padx=4,pady=4,font=self.heading,anchor="center")
        self.labelv.grid(row=0,column=1,columnspan=4, sticky=S)
        
        self.vStart=1.5
        self.vLower=Button(self.sm,text="<",fg="black",bg="gray",height=2,width=7,command=self.vdecrease)
        self.vLower.grid(row=1,column=1)
        self.labelvStart=Label(self.sm,text=self.vStart,padx=4,pady=4,font=self.content,anchor="center")
        self.labelvStart.grid(row=1,column=2,columnspan=2,sticky=N+E+S+W)
        self.vUpper=Button(self.sm,text=">",fg="black",bg="gray",height=2,width=7,command=self.vincrease)
        self.vUpper.grid(row=1,column=4)
        
        self.vSet=Button(self.sm,text="Set Total Volume",height=2,width=14,command=self.vset)
        self.vSet.grid(row=2,column=1,columnspan=2,sticky=W)
        self.vReset=Button(self.sm,text="Reset to Default",height=2,width=14,command=self.vdefault)
        self.vReset.grid(row=2,column=3,columnspan=2,sticky=E)
        
        
        ##Cycle Control
        self.labelc=Label(self.sm,text="Cycles Control",padx=4,pady=2,font=self.heading,anchor="center")
        self.labelc.grid(row=3,column=1,columnspan=4,sticky=S)
        
        self.cStart=20
        self.cLower=Button(self.sm,text="<",fg="black",bg="gray",height=2,width=7,command=self.cdecrease)
        self.cLower.grid(row=4,column=1)
        self.labelcStart=Label(self.sm,text=self.cStart,padx=4,pady=4,width=14,font=self.content,anchor="center")
        self.labelcStart.grid(row=4,column=2,columnspan=2,sticky=N+E+W+S)
        self.cUpper=Button(self.sm,text=">",fg="black",bg="gray",height=2,width=7,command=self.cincrease)
        self.cUpper.grid(row=4,column=4,sticky=E)

        self.cSet=Button(self.sm,text="Set Cycle Number",height=2,width=14,command=self.cset)
        self.cSet.grid(row=5,column=1,columnspan=2,sticky=W)
        self.cReset=Button(self.sm,text="Reset to Default",height=2,width=14,command=self.cdefault)
        self.cReset.grid(row=5,column=3,columnspan=2,sticky=E)
        
        
        #Output Information
        self.statusLabel=Label(self.sm,text="Status Overview",padx=4,pady=4,font=self.heading,anchor="center")
        self.statusLabel.grid(row=0,column=5,columnspan=2,sticky=E+S+W)
        
        self.cycleCount=0
        self.cCounterLabel=Label(self.sm,text="Current Cycle",padx=4,pady=4,width=31,font=self.content,anchor="center")
        self.cCounterLabel.grid(row=0,column=5,columnspan=2,sticky=E+S+W)
        self.cycleCountLabel=Label(self.sm,text=self.cycleCount,padx=4,pady=4,width=31,font=self.content,anchor="center")
        self.cycleCountLabel.grid(row=1,column=5,columnspan=2,sticky=N+E+W)
        
        self.now="00:00:00"
        self.timeElapsedLabel=Label(self.sm,text="Time Elapsed",padx=4,pady=4,font=self.content,anchor="center")
        self.timeElapsedLabel.grid(row=2,column=5,sticky=S+E+W)
        self.timeUp=Label(self.sm,text=self.now,padx=4,pady=4,font=self.content,anchor="center")
        self.timeUp.grid(row=3,column=5,sticky=N+E+W)
        
        self.remain="00:00:00"
        self.timeRemainLabel=Label(self.sm,text="Time Remaining",padx=4,pady=4,font=self.content,anchor="center")
        self.timeRemainLabel.grid(row=2,column=6,sticky=S+E+W)
        self.timeDown=Label(self.sm,text=self.remain,padx=4,pady=4,font=self.content,anchor="center")
        self.timeDown.grid(row=3,column=6,sticky=N+E+W)

        #Output text widget
        self.text_box=Text(self.sm,padx=4,pady=4,height=6,width=34,state=DISABLED)
        self.text_box.grid(row=4,column=5,columnspan=2,rowspan=2)
        self.scrollb = Scrollbar(self.sm,command=self.text_box.yview)
        self.scrollb.grid(row=4, column=6,rowspan=2,sticky=N+S+E)
        self.text_box['yscrollcommand'] = self.scrollb.set
        
        sys.stdout = StdRedirector(self.text_box)
        sys.stderr = StdRedirector(self.text_box)
        
        #Parameters
        stat=MOTOR.getSENSORS(0)
        self.homeComplete=0
        self.lysing=0
        self.vFinal=0
        self.vCurrent=0
        self.cFinal=0
        self.direction=0
        self.dir_dict={0:'cw', 1:'ccw'}
        
        
            
        GPIO.setmode(GPIO.BCM)  
        GPIO.setup(22, GPIO.IN, pull_up_down=GPIO.PUD_UP)  #set up GPIO 22 as an input
        MOTOR.enablestepSTOPint(0,'a')
        MOTOR.enablestepSTOPint(0,'b')
        MOTOR.intEnable(0)
        
        
    #Change volume functions
    def vset(self):
        if self.homeComplete:
            self.vFinal=self.vStart
            print "Moving syringe A to",self.vFinal,"mL"
        else:
            print "Please home motors first"
        
    def vdefault(self):
        if self.homeComplete:
            #Set volume back to default value of 1.5mL
            self.vStart=1.5
            self.labelvStart.config(text=self.vStart)
        else:
            print "Please home motors first"
    
    def vdecrease(self):
        #Decrease volume by 0.1mL to a minimum of 0.5mL
        if self.homeComplete:
            if self.vStart > 0.5:
                self.vStart-=0.1
                self.vStart = round(self.vStart,1)
                self.labelvStart.config(text=self.vStart)
                #print(self.vStart)
        else:
            print "Please home motors first"
            
    def vincrease(self):
        #Increase the volumeby 0.1mL to a maximum of 3.0mL
        if self.homeComplete:
            if self.vStart < 3:
                self.vStart+=0.1
                self.vStart = round(self.vStart,1)
                self.labelvStart.config(text=self.vStart)
                #print(self.vStart)
        else:
            print "Please home motors first"
    
    #Change cycles functions
    def cset(self):
        if self.homeComplete:
            self.cFinal=self.cStart
            print "Set cycle number to",self.cFinal
        else:
            print "Please home motors first"
        
    def cdefault(self):
        if self.home.Complete:
            #Reset cycle number to a default value of 20
            self.cStart=20
            self.labelcStart.config(text=self.cStart)  
        else:
            print "Please home motors first"
            
    def cdecrease(self):
        if self.homeComplete:
            #Decrease cycle number by 1
            if self.cStart > 1:
                self.cStart-=1
                self.labelcStart.config(text=self.cStart)
                #print(self.vStart)
        else:
            print "Please home motors first"
    
    def cincrease(self):
        if self.homeComplete:            
            #Increase cycle number by 1
            self.cStart+=1
            self.labelcStart.config(text=self.cStart)
            #print(self.vStart)
        else:
            print "Please home motors first"
            
            
    #Home functions
    def jogHome(self):
        #Initialize homing variables
        self.motorA=1
        self.motorB=1
        #Configure motor parameters for jogging
        self.sync_config(1,150,0)
        #Move steppers A and B towards switches
        MOTOR.stepperJOG(0,'a')
        MOTOR.stepperJOG(0,'b')
        #Wait for limit switch signal
        root.after(1,self.findHome)

    def findHome(self):
        #Sample every 100ms
        self._job=root.after(100,self.findHome)
        self.stat=MOTOR.getSENSORS(0)
        #If limit switch A is hit, stop and reverse
        if not (self.stat&0x01):
            MOTOR.stepperSTOP(0,'a')
            MOTOR.stepperCONFIG(0,'a','cw','M8',100,0)
            MOTOR.stepperMOVE(0,'a',50)
            MOTOR.stepperOFF(0,'a')
            self.motorA=0
            print "MOTOR A homing complete"
        #If limit switch B is hit, stop and reverse
        if not (self.stat&0x04):
            MOTOR.stepperSTOP(0,'b')
            MOTOR.stepperCONFIG(0,'b','cw','M8',100,0)
            MOTOR.stepperMOVE(0,'b',50)
            MOTOR.stepperOFF(0,'b')
            self.motorB=0
            print "MOTOR B homing complete"
        #When both motors are stopped, cancel after function
        if (self.motorA==0 and self.motorB==0):
            self.vCurrent=0
            self.homeComplete=1
            print"Home complete"
            root.after_cancel(self._job)
      
    #Lysate making functions  
    def makeLysate(self):
        global intBITS, intFLAG
        self.lysing=1
        self.cycleCount=0
        print self.cFinal
        

        if self.homeComplete:
            GPIO.setmode(GPIO.BCM)  
            GPIO.setup(22, GPIO.IN, pull_up_down=GPIO.PUD_UP)  #set up GPIO 22 as an input 
            GPIO.add_event_detect(22, GPIO.FALLING, callback=self.moveDone)  # detect interrupts
            MOTOR.enablestepSTOPint(0,'a')
            MOTOR.enablestepSTOPint(0,'b')
            MOTOR.intEnable(0)
            print "Lysing"
            print self.cycleCount
            self.sync_config(self.direction,120,0)
            root.after(1,self.movement)        
        else:
            print("Remove syringes and home motors")
    
    def movement(self):
        global intBITS, intFLAG
        if self.cycleCount < 5:
            print self.cycleCount
            self.sync_move(500)
            print "moving"
            print intBITS
            print intFLAG
            root.after(100,self.doCycle)
            root.after(100,self.limitSwitch)
        else:
            print "No more"
            #GPIO.remove_event_detect(22, GPIO.FALLING, callback=self.moveDone) 
                 
    def doCycle(self):
        global intBITS, intFLAG
        if intFLAG:
            print "intflag"
            intFLAG=0
            if not intBITS&0x20:
                print "stopping"
                self.direction = self.toggle_direction(self.direction)
                self.sync_config(self.direction,120,0)
                print "changing directions"
                self.cycleCount +=0.5
                print self.cycleCount
                
        self._job=root.after(500,self.movement)
    
    def toggle_direction(self,dir_val):
        return abs(dir_val-1)
    
    def sync_config(self,dir,rate,acc):
        MOTOR.stepperCONFIG(0,'a',self.dir_dict[dir],'M8',rate,acc)
        MOTOR.stepperCONFIG(0,'b',self.dir_dict[self.toggle_direction(dir)],'M8',rate,acc)
    
    def sync_move(self,steps):
        MOTOR.stepperMOVE(0,'a',steps)
        MOTOR.stepperMOVE(0,'b',steps)
    
    def sync_stop(self):
        MOTOR.stepperSTOP(0,'a')
        MOTOR.stepperSTOP(0,'b')
    
    #Interrupt functions
    def moveDone(self, channel):
        # signal to main program that an int has occurred
        global intBITS, intFLAG
        intBITS=MOTOR.getINTflag0(0)
        intFLAG=1
        
    def limitSwitch(self):
        self.stat=MOTOR.getSENSORS(0)
        if not (self.stat&0x01) or not (self.stat&0x04):
            print "Limit switches hit, stopping"
            self.stopAll
    
    #Stopping functions
    def stopAll(self):
        #root.after_cancel(self._job)
        MOTOR.RESET(0)
        self.homeComplete=0
        self.lysing=0
        self.vFinal=0
        self.cFinal=0
        self.cycleCount=0
        self.intReset
        print("Stopping and resetting all processes")

    def intReset(self):
        global intBITS, intFLAG
        intBITS=MOTOR.getINTflag0(0)
        intFlag=1
        GPIO.remove_event_detect(22)
        GPIO.cleanup() 


    #Clock
    def update_clock(self):
        self.now = time.strftime("%H:%M:%S")
        self.timeUp.configure(text=self.now)
        self.root.after(1000, self.update_clock)
        

   
class StdRedirector():
    def __init__(self, text_widget):
        self.text_space = text_widget

    def write(self, string):
        self.text_space.config(state=NORMAL)
        self.text_space.insert("end", string)
        self.text_space.see("end")
        self.text_space.config(state=DISABLED)

MOTOR.RESET(0)  
        
root = Tk()
root.config(bg="black")
root.attributes("-fullscreen", TRUE)
swidth=root.winfo_screenwidth()
sheight=root.winfo_screenheight()
#print tkFont.families()
container=Frame(root,bg="white")
container.place(relx=0.5, rely=0.5, anchor=CENTER)

title = headerBLOCK(container,0,0)
control = controlGUI(container,1,0)

root.mainloop()
```
