## Supplementary material for "Low-Cost Touchscreen Driven Programmable Dual Syringe Pump for Life Science Applications": SyncMvmt_JL2.py

```
import piplates.MOTORplate as MOTOR
import RPi.GPIO as GPIO
import time
import os
import string
from Tkinter import *
import tkFont


class controlGUI:
    global intBITS, intFLAG
    intBITS=MOTOR.getINTflag0(0)
    intFLAG=0
    

    
    

    def __init__(self,master,r,c):
    
        self.master=master
        self.initialization=False
        self.sm=Frame(self.master,padx=4,pady=4,bd=2,relief='sunken')
        self.sm.grid(row=r,column=c, sticky=N+S+W+E)
        
        ##Create Fonts
        self.title=tkFont.Font(family='Helvetica', size=25, weight='bold') 
        self.heading=tkFont.Font(family='Helvetica', size=18, weight='bold')
        self.content=tkFont.Font(family='Helvetica', size=12, weight='bold')


        self.dir_dict = {0:'cw', 1:'ccw'}
        self.direction=1
        self.cycleCount=1
        
        self.bigStart=Button(self.sm,text="go",height=108,width=125,command=self.makeLysate)
        self.bigStart.grid(row=2,column=0,rowspan=2,padx=30,pady=4)
        #self.dummy = MOTOR.getINTflag0(0)
        

        
        

#allow interrupts when a motor stops
    def intReset(self):
        global intBITS, intFLAG
        intBITS=MOTOR.getINTflag0(0)
        intFlag=0
        GPIO.cleanup() 
    

    
    def moveDone(self, channel):
        # signal to main program that an int has occurred
        global intBITS, intFLAG
        intBITS=MOTOR.getINTflag0(0)
        intFLAG=1
        

    
    def makeLysate(self):
        global intBITS, intFLAG
        GPIO.setmode(GPIO.BCM)  
        GPIO.setup(22, GPIO.IN, pull_up_down=GPIO.PUD_UP)  #set up GPIO 22 as an input 
        GPIO.add_event_detect(22, GPIO.FALLING, callback=self.moveDone)  # detect interrupts
        MOTOR.enablestepSTOPint(0,'a')
        MOTOR.enablestepSTOPint(0,'b')
        MOTOR.intEnable(0)
        print "Lysing"
        print self.cycleCount
        self.sync_config(self.direction,120,0)
        root.after(1,self.movement)

    def movement(self):
        global intBITS, intFLAG
        if self.cycleCount < 3:
            self.sync_move(500)
            print "moving"
            print intBITS
            print intFLAG
            root.after(100,self.doCycle)
            root.after(100,self.checkint)
        else:
            print "No more"
    
    def checkint(self):
        self.stat=MOTOR.getSENSORS(0)
        if not self.stat&0x04:
            MOTOR.RESET(0)
        elif not self.stat&0x01:
            MOTOR.RESET(0)
             
    def doCycle(self):
        global intBITS, intFLAG
        if intFLAG:
            print "int flag"
            print intBITS&0x20
            intFLAG=0
            if not intBITS&0x20:
                print "stopping"
                self.direction = self.toggle_direction(self.direction)
                self.sync_config(self.direction,120,0)
                print "changing directions"
                #self.sync_move(500)
                self.cycleCount +=1
                print self.cycleCount
        self._job=root.after(100,self.movement)
    
            
            
            

    
    def limitSwitch(self):
        self.stat=MOTOR.getSENSORS(0)
        if not (self.stat&0x01) or not (self.stat&0x04):
            print "Limit switches hit, stopping"
            self.stopAll
    
    def toggle_direction(self,dir_val):
        return abs(dir_val-1)
    
    def sync_config(self,dir,rate,acc):
        MOTOR.stepperCONFIG(0,'a',self.dir_dict[dir],'M8',rate,acc)
        MOTOR.stepperCONFIG(0,'b',self.dir_dict[self.toggle_direction(dir)],'M8',rate,acc)
    
    def sync_move(self,steps):
        MOTOR.stepperMOVE(0,'a',steps)
        MOTOR.stepperMOVE(0,'b',steps)
    
    def sync_stop(self):
        MOTOR.stepperSTOP(0,'a')
        MOTOR.stepperSTOP(0,'b')
    


root = Tk()
root.config(bg="black")
root.attributes("-fullscreen", FALSE)
swidth=root.winfo_screenwidth()
sheight=root.winfo_screenheight()
#print tkFont.families()
container=Frame(root,bg="white")
container.place(relx=0.5, rely=0.5, anchor=CENTER)

control = controlGUI(container,0,0)

root.mainloop()
```
