## Supplementary material for "Low-Cost Touchscreen Driven Programmable Dual Syringe Pump for Life Science Applications": SyncMvmt_JL.py

```
import piplates.MOTORplate as MOTOR
import RPi.GPIO as GPIO
import time
import os
import string
from Tkinter import *
import tkFont


#flush out any old interrupts
dummy = MOTOR.getINTflag0(0)
intFLAG=0
intITS=0

def moveDone(channel):
    global intBITS, intFLAG
    intBITS=MOTOR.getINTflag0(0)
    intFLAG=1

MOTOR.RESET(0)


dir_dict = {0:'cw', 1:'ccw'}

def toggle_direction(dir_val):
    return abs(dir_val-1)
    
def sync_config(dir,rate,acc):
    MOTOR.stepperCONFIG(0,'a',dir_dict[dir],'M8',rate,acc)
    MOTOR.stepperCONFIG(0,'b',dir_dict[toggle_direction(dir)],'M8',rate,acc)
    
def sync_move(steps):
    MOTOR.stepperMOVE(0,'a',steps)
    MOTOR.stepperMOVE(0,'b',steps)
    
def sync_jog():
    MOTOR.stepperJOG(0,'a')
    MOTOR.stepperJOG(0,'b')
    
def sync_stop():
    MOTOR.stepperSTOP(0,'a')
    MOTOR.stepperSTOP(0,'b')
    
    
def make_lysate(cycles,steps,direction):
    counter=0
    while (counter<=cycles):
        sync_move(steps)
        direction=toggle_direction(direction)
        time.sleep(1)
        sync_config(direction,500,1)
        sync_move(1500)
        counter+=1
        
        
#GPIO.setmode(GPIO.BCM)
#GPIO.setup(22, GPIO.IN, pull_up_down=GPIO.PUD_UP)
#GPIO.add_event_detect(22, GPIO.FALLING, callback=moveDone)
#allow interrupts when a motor stops
#MOTOR.enablestepSTOPint(0,'a')
#MOTOR.enablestepSTOPint(0,'b')
#MOTOR.intEnable(0)
        
print MOTOR.getID(1)  
 
dummy=MOTOR.getINTflag0(1)  #flush out any old interrupts
intFLAG=0
intITS=0

direction=1

sync_config(direction,200,0)
#sync_jog()

flag=0
counter=0
cycles=0
sync_move(1)
while (counter<cycles):
    time.sleep(0.1)
    stat=MOTOR.getSENSORS(0)
    intBITS=MOTOR.getINTflag0(0)
    sync_move(2)
    if(intBITS&0x20):
        time.sleep(3)
        direction=toggle_direction(direction)
        sync_config(direction,100,1)
        print (direction)
        print (counter)
        counter+=1
        sync_move(1000)
    if not (stat&0x01) or not (stat&0x04):
        sync_stop()
        direction=toggle_direction(direction)
        time.sleep(2)
        sync_config(direction,100,1)
        sync_move(1000)
    if not (stat&0x01) and not (stat&0x04):
        sync_stop()
#	MOTOR.RESET(0)
#	sys.exit()
class controlGUI:
    def __init__(self,master,r,c):
        self.master=master
        self.initialization=False
        self.sm=Frame(self.master,padx=4,pady=4,bd=2,relief='sunken')
        self.sm.grid(row=r,column=c, sticky=N+S+W+E)
        
        ##Create Fonts
        self.title=tkFont.Font(family='Helvetica', size=25, weight='bold') 
        self.heading=tkFont.Font(family='Helvetica', size=18, weight='bold')
        self.content=tkFont.Font(family='Helvetica', size=12, weight='bold')


        self.dir_dict = {0:'cw', 1:'ccw'}
        self.direction=1
        self.cycleCount=1
        
        self.bigStart=Button(self.sm,text="go",height=108,width=125,command=self.makeLysate)
        self.bigStart.grid(row=2,column=0,rowspan=2,padx=30,pady=4)
        self.intBITS=MOTOR.getINTflag0(0)
        self.intFLAG=1
        
                
        GPIO.setmode(GPIO.BCM)
        GPIO.setup(22, GPIO.IN, pull_up_down=GPIO.PUD_UP)
        GPIO.add_event_detect(22, GPIO.FALLING, callback=self.moveDone)
#allow interrupts when a motor stops
        MOTOR.enablestepSTOPint(0,'a')
        MOTOR.enablestepSTOPint(0,'b')
        MOTOR.intEnable(0)
        
    def moveDone(self):
        global intBITS, intFLAG
        self.intBITS=MOTOR.getINTflag0(0)
        self.intFLAG=1
        
    def makeLysate(self):
        print "Lysing"
        print self.cycleCount
        #self.sync_config(self.direction,120,0)
        if self.cycleCount <=3:
            #sync_move(500)
            #print "moving"
            self.moveDone
            print self.intBITS
            root.after(100,self.doCycle)
            root.after(100,self.checkint)
    
    def checkint(self):
        self.stat=MOTOR.getSENSORS(0)
        if not (self.stat&0x04 or self.stat&0x01):
            MOTOR.reset(0)
             
    def doCycle(self):

        if self.intFLAG:
            self.intFLAG=0
            if self.intBITS&0x20:
                print "stopping"
                self.direction = toggle_direction(self.direction)
                self.sync_config(self.direction,120,0)
                print "changing directions"
                sync_move(500)
                self.cycleCount +=1
                print self.cycleCount
        self._job=root.after(100,self.makeLysate)
    
            
            
            

    
    def limitSwitch(self):
        self.stat=MOTOR.getSENSORS(0)
        if not (self.stat&0x01) or not (self.stat&0x04):
            print "Limit switches hit, stopping"
            self.stopAll
    
    def toggle_direction(self,dir_val):
        return abs(dir_val-1)
    
    def sync_config(self,dir,rate,acc):
        MOTOR.stepperCONFIG(0,'a',self.dir_dict[dir],'M8',rate,acc)
        MOTOR.stepperCONFIG(0,'b',self.dir_dict[self.toggle_direction(dir)],'M8',rate,acc)
    
    def sync_move(self,steps):
        MOTOR.stepperMOVE(0,'a',steps)
        MOTOR.stepperMOVE(0,'b',steps)
    
    def sync_stop(self):
        MOTOR.stepperSTOP(0,'a')
        MOTOR.stepperSTOP(0,'b')
    


root = Tk()
root.config(bg="black")
root.attributes("-fullscreen", FALSE)
swidth=root.winfo_screenwidth()
sheight=root.winfo_screenheight()
#print tkFont.families()
container=Frame(root,bg="white")
container.place(relx=0.5, rely=0.5, anchor=CENTER)

control = controlGUI(container,0,0)

root.mainloop()
```
