## Supplementary material for "Low-Cost Touchscreen Driven Programmable Dual Syringe Pump for Life Science Applications": test_Move_JL.py

```
import piplates.MOTORplate as MOTOR
import RPi.GPIO as GPIO
import time
import os
import string
from Tkinter import *
import tkFont

print MOTOR.getID(0)

MOTOR.RESET(0)

#configure steppers A and B for homing function
MOTOR.stepperCONFIG(0,'a','cw','M8',100,0)
MOTOR.stepperCONFIG(0,'b','ccw','M8',500,0)


MOTOR.stepperMOVE(0,'a',7100)
```
