## Supplementary figures and images for "Low-Cost Touchscreen Driven Programmable Dual Syringe Pump for Life Science Applications"

### 3D-ppLogo.gif

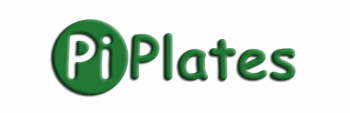

### 2018-02-14-164246_800x480_scrot.png

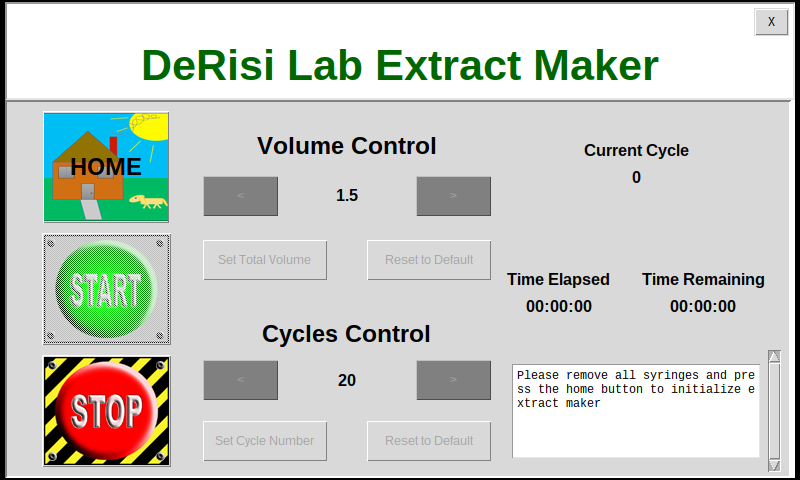

### easteregg.jpg

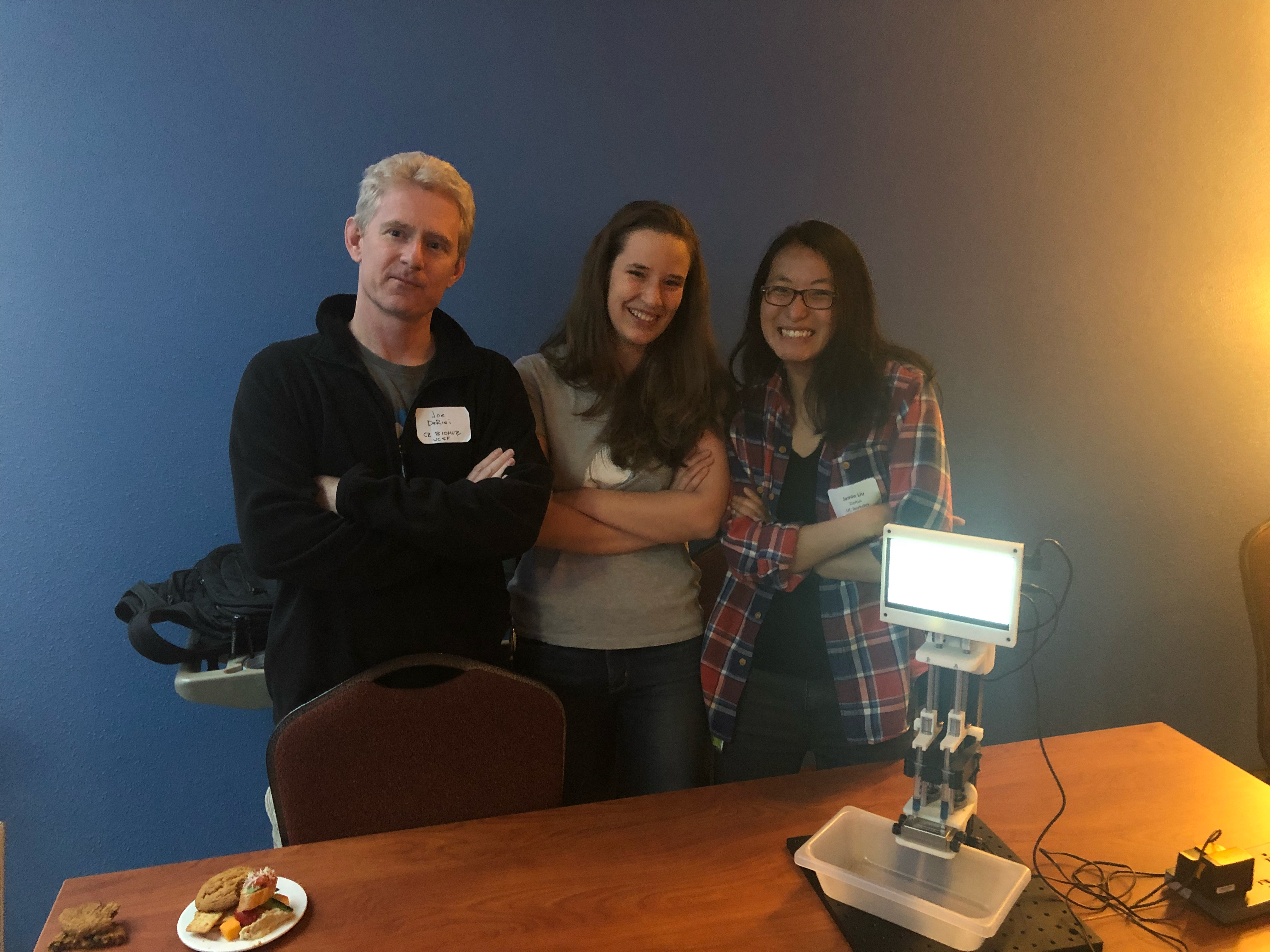

### StartPushSmall.png

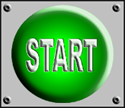

### StopPushSmall.png

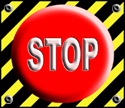

### wes_home.png

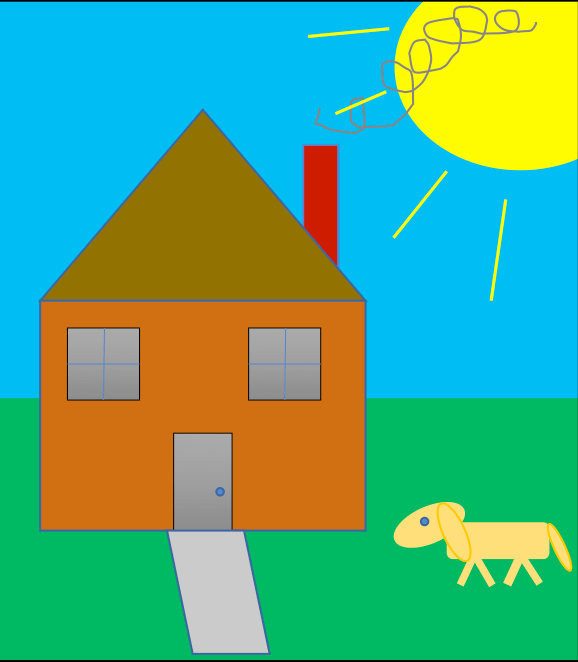

### wes_home_orig.png

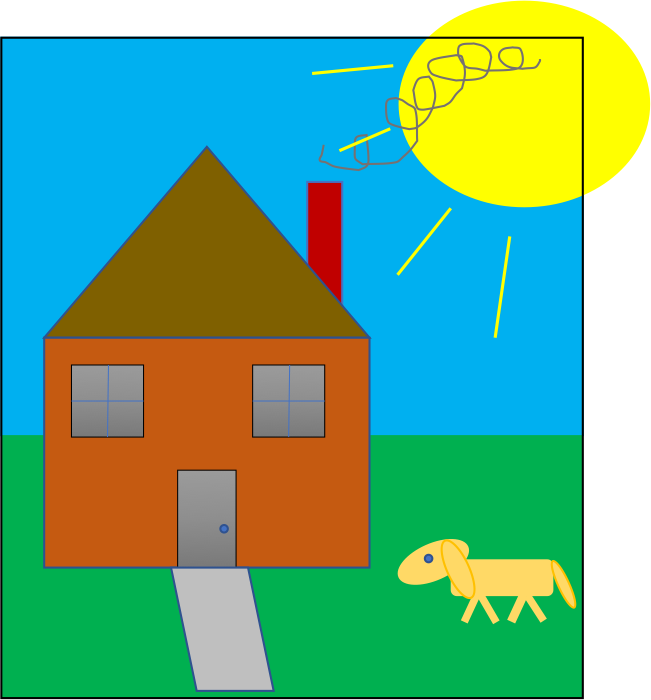
